# Rhizobia transform legume nodules via native T-DNA transfer systems

**DOI:** 10.64898/2026.11.03.763973

**Authors:** Shan-Chi Hsieh, Erh-Min Lai, Chih-Horng Kuo

**Affiliations:** Institute of Plant and Microbial Biology, Academia Sinica, Taiwan

**Keywords:** rhizobia, plant transformation, T-DNA, type IV secretion system, root nodule, agrobacteria

## Abstract

Agrobacteria can deliver transfer DNA (T-DNA) into plant cells, causing tumorous growth. This process, agrobacteria-mediated transformation (AMT), underlies plant genetic engineering and has been regarded as an evolutionary innovation specific to these pathogens. By screening 63,759 alphaproteobacterial genomes for T-DNA border sequences, we identify mini-T-DNAs in diverse rhizobial genera. Each typically encodes a single cargo, a RolB/RolC family protein or a distant rhizobial homologue. We show that native rhizobia-mediated transformation (RMT) systems deliver these mini-T-DNAs into the root nodules of common bean, pea, and chickpea, where the cargo genes are expressed, with no signal in adjacent root tissue. RMT systems share VirG-dependent activation of the core virulence regulon but respond to distinct upstream signals. Our genome screen identifies many additional candidate T-DNA transfer systems, and phylogenetic analysis places their genes into two of the many type IV secretion system (T4SS) clades. RMT systems occur in both clades, together with other systems unrelated to T-DNA transfer, and typically have an ancestral-like T4SS configuration. In contrast, AMT systems occupy a single subclade, consistent with later acquisition of their T4SS module. AMT and RMT thus represent two specialized outcomes of an older, more widespread capability for bacteria-to-plant DNA transfer.

## Introduction

Bacteria-to-plant DNA transfer is a rare evolutionary innovation with profound consequences for both plant biology research and biotechnology applications. The well-characterized system of agrobacteria-mediated transformation (AMT), in which oncogenic plasmids encode the machinery for delivering defined DNA segments, termed transfer DNA (T-DNA), into plant cells for transformation. These plasmids fall into two major classes: tumor-inducing plasmids (pTi)^1^ and root-inducing plasmids (pRi)^2^, named for the tumorous crown gall and proliferative hairy root tissues they induce, respectively. Despite producing distinct disease phenotypes, both plasmid classes share a conserved T-DNA transfer module: border repeats that delimit the transferred DNA segment and virulence (*vir*) genes encoding functional components for transformation. Key Vir proteins include VirD2, the relaxase that nicks and remains attached to the T-DNA strand; VirE proteins that protect the T-DNA and facilitate its delivery to the nucleus^3–6;^ and the VirB apparatus and VirD4 coupling protein that assemble into the type IV secretion system (T4SS), which transports the T-DNA and associated effectors into plant cells. After delivery, T-DNA cargos typically encode opine biosynthesis genes together with oncogenic determinants that manipulate host physiology. The best-characterized examples are the auxin and cytokinin biosynthesis enzymes of pTi^7–10^, and the RolB and RolC proteins of pRi. These belong to the RolB/RolC family, defined as containing the Pfam PF02027 domain, and correspond to the phenotypic-plasticity (plast) family of agrobacterial T-DNA oncoproteins^11^; for brevity, we refer to all PF02027-containing proteins collectively as RolB hereafter. These proteins alter auxin sensitivity, promote adventitious root and meristem formation^12–15^, and exhibit diverse and sometimes antagonistic activities among individual members^16^.

The bacteria that carry these oncogenic plasmids, collectively termed agrobacteria, have emerged repeatedly within the agrobacteria-rhizobia complex (ARC)^17^, a broader grouping that also includes nitrogen-fixing, root-nodulating rhizobia and other plant-associated bacteria. This polyphyletic distribution of agrobacterial lineages reflects the mobility of diverse oncogenic plasmids and underscores the importance of plasmid-level rather than organism-level thinking when considering the evolutionary origins of AMT. The modular organization of these plasmids, in which T-DNA, transfer machinery, and regulatory components can be exchanged independently, has facilitated their diversification and horizontal dissemination across the ARC^18,19^. Yet a clear discrepancy exists: the core genetic components required for T-DNA transfer are distributed far more broadly across the ARC than the oncogenic plasmids. This raises three connected questions. First, at the plasmid level: do homologous systems outside canonical oncogenic plasmids retain functional T-DNA transfer capacity? Second, at the taxon level: if so, is this capacity distributed beyond known agrobacteria? Third, at the evolutionary level: if non-oncogenic T-DNA transfer systems exist, do they shed light on the ancestral form of this machinery, and on how AMT emerged as a specialized pathogenic innovation?

Addressing these questions has been hampered by a fundamental ascertainment bias. T-DNA transfer has historically been inferred from disease phenotypes, an operationally convenient but conceptually narrow criterion that systematically excludes non-oncogenic systems. As a result, AMT-like systems outside canonical oncogenic plasmids have generally been interpreted either as degraded remnants of pathogenic systems or as machinery repurposed for other functions. The former interpretation is illustrated by “partial virulence” plasmids, including the Ti-like plasmid p42a of *Rhizobium etli* CFN42, which are implicated in T-DNA transfer without causing disease^19^. The latter is illustrated by the T4SS encoded by the ICESym element of *Mesorhizobium japonicum* R7A^20^, which translocates symbiosis-promoting proteins rather than T-DNA into host cells^21^. Consequently, the possibility that intact, non-oncogenic T-DNA transfer systems exist has not been systematically investigated.

To overcome this bias and address the three questions above, we combine large-scale comparative genomics with functional validation, enabling systematic identification of T-DNA transfer systems independent of disease phenotype. By screening 63,759 alphaproteobacterial genomes for T-DNA border sequences, we identify a class of short rhizobial T-DNAs that typically encode a single RolB or RolB-like cargo protein. We show that multiple rhizobial strains can mediate transfer of these T-DNAs into root nodules of their native host plants, defining a class of rhizobia-mediated transformation (RMT) systems. Phylogenetic analyses of the T4SS components reveal that many RMT systems are associated with a distinct, ancestral-like configuration of the transfer machinery, suggesting that the T4SS module used by canonical AMT systems was acquired later in a modular replacement event. Together, these findings reframe bacteria-to-plant DNA transfer from a uniquely agrobacterial pathogenic innovation to a broader evolutionary strategy among plant-associated bacteria, with AMT and RMT representing two specialized subclasses of an ancient and widely distributed system.

## Results

### Bioinformatic screening reveals rhizobial mini-T-DNAs encoding RolB/RolB-like proteins

We screened 63,759 alphaproteobacterial genomes for T-DNA border sequences matching the consensus motif (YGRCAGGATATATNNNNNKGTMAWN)^22^, identifying 1,706 candidate T-DNAs, and revealed an unexpected enrichment of compact T-DNA loci in rhizobia. To reduce false positives, we applied additional criteria based on features of known T-DNAs.

Previous studies have shown that T-DNAs typically exhibit lower coding sequence (CDS) density and GC content^23^. Using loci containing the conserved auxin biosynthesis gene, *tms1* (*rolB*-*iaaM*), present in most known pTi and pRi as an empirical positive-control set^18^, we defined likely T-DNAs as loci with <80% CDS density and a GC content more than 7% lower than that of the host genome (Fig. 1a). Consistent with the notion that most *vir* gene homologues were concentrated within Hyphomicrobiales (synonym Rhizobiales), nearly all candidate T-DNAs identified outside this order were removed by these filters, thus were considered as false positives.

**Fig. 1.**
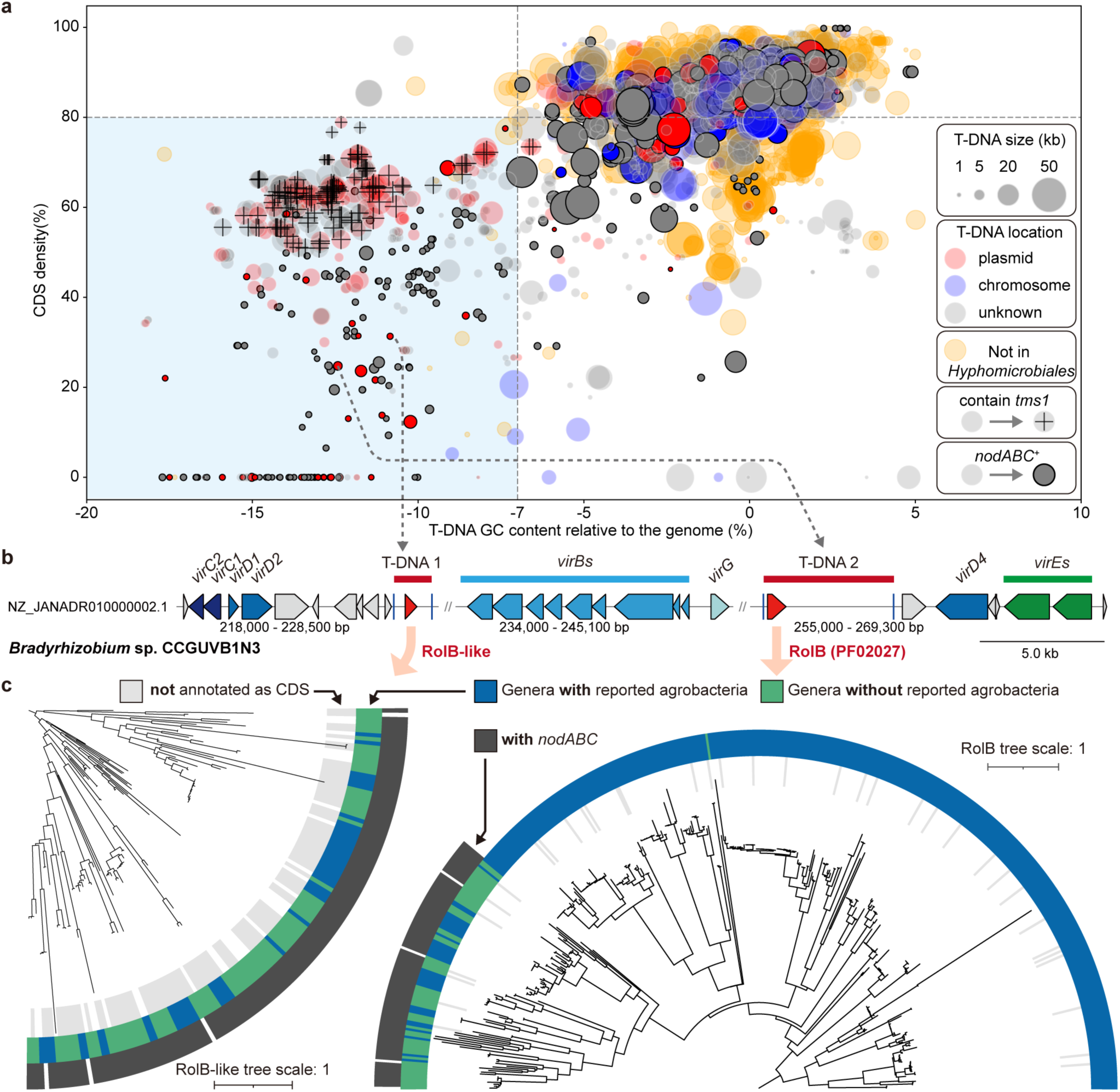
Identification of mini-T-DNAs carrying *rolB* and novel *rolB*-like genes in rhizobia by a genome-wide T-DNA search. **a**, T-DNAs predicted from flanking border-like sequences, plotted by coding-sequence (CDS) density against GC content relative to the host genome. Dot size indicates T-DNA length; dot colour denotes genomic location (plasmid, chromosome, or unknown) and whether the sequence originates from taxa outside the Hyphomicrobiales; a cross marks the presence of the conserved agrobacterial *tms1* gene as cargo; and dot transparency indicates the presence or absence of a *nodABC* operon in the corresponding genome. The blue shaded region highlights candidates considered likely to represent bona fide T-DNAs. **b**, Two T-DNAs from a *nodABC*⁺ genome, with homologues of *Agrobacterium* T-DNA transfer genes indicated. One T-DNA carries a *rolB* family gene; the other encodes a gene of unknown function that shows similarity to the *rolB* family in profile-to-profile searches (*rolB*-like). **c**, Phylogenetic trees of RolB and RolB-like proteins identified in the Hyphomicrobiales. Colour strips indicate the host genus, the presence or absence of a *nodABC* operon in the host genome, and whether each sequence was annotated as a coding sequence in the source genome. Trees are midpoint-rooted.

Notably, many likely T-DNAs were compact loci occurring outside the six genera known to include agrobacteria. Most occurred in genomes carrying the *nodABC* gene cluster, which we use as a genomic marker for rhizobia, the nitrogen-fixing symbionts of legume root nodules (Fig. 1a). We refer to these compact, single-cargo loci as mini-T-DNAs. We identified 157 such loci, of which 80 contain a *rolB* family member, 12 encode short ORFs with no detectable homologues, and 54 lack annotated coding sequences. For example, *Bradyrhizobium* sp.

CCGUVB1N3 contains two mini-T-DNAs adjacent to *vir* genes: one encoding a RolB family protein and the other a small ORF of unknown function (Fig. 1b). Although hidden Markov model (HMM) searches did not detect homology between this ORF and RolB family proteins, the more sensitive HHsearch identified significant similarity, suggesting that it encodes a RolB-like protein.

We therefore constructed a new HMM profile using 44 RolB-like homologues identified by RPS-BLAST. Scanning six-frame translations of alphaproteobacterial genomes revealed that most apparently empty mini-T-DNAs in fact encode previously unannotated *rolB*-like genes (53/54). Despite their small size (100-150 amino acids), structural predictions indicate that these proteins are not simply N-terminal or C-terminal truncated RolB variants, but instead adopt a fold resembling the core structure of RolB family proteins (Extended Data Fig. 1).

To assess their distribution, we constructed phylogenetic trees of RolB and RolB-like proteins, annotated by host taxonomy and *nodABC* presence. The newly identified *rolB*-like genes were predominantly found in rhizobia, with a *nodABC* cluster detected in nearly all host genomes (129/133). By contrast, *rolB* occurred in both agrobacterial and rhizobial lineages, with closely related sequences distributed across multiple genera, consistent with frequent horizontal gene transfer, as expected given that most *rolB* and *rolB*-like genes are plasmid-encoded. Despite this mobility, the rhizobial RolB proteins formed a distinct clade, consistent with lineage-specific diversification and possible selective constraints (Fig. 1c). Together, these analyses identify a subset of rhizobia with specialized T-DNA transfer systems: compact T-DNAs carrying divergent *rolB*/*rolB*-like cargos occur in approximately 2% of genomes containing a *nodABC* cluster.

### Rhizobial mini-T-DNAs retain canonical border features but differ in cargo organization

Agrobacterial T-DNAs often contain multiple *rolB* genes, encoding proteins of ∼200-300 amino acids, as well as a conserved *rolB-iaaM* fusion gene (*tms1*; encodes ∼750 amino acids) present in most known pTi/pRi^15,18^, resulting in variable distances between *rolB* and T-DNA borders. This variability is illustrated by aligning non-redundant *rolB* genes at their stop codons and plotting the relative positions of borders and start codons (Fig. 2a, left). By contrast, analogous plots for rhizobial *rolB* and *rolB*-like genes revealed two pronounced border-position peaks on the opposite DNA strand, located close to the start and stop codons (Fig. 2a). This pattern is consistent with rhizobial *rolB/rolB*-like genes typically occurring as single cargos within compact T-DNAs and being oriented opposite to the border sequences, unlike agrobacterial T-DNAs, in which *rolB* genes are often co-oriented with the borders. Supporting their assignment as genuine T-DNAs, the borders flanking rhizobial *rolB/rolB*-like genes displayed canonical features of agrobacterial T-DNAs: left borders were relatively A-rich at their 3′ ends, whereas right borders frequently contained a downstream T-rich “overdrive” sequence (Fig. 2b)^24^.

**Fig. 2.**
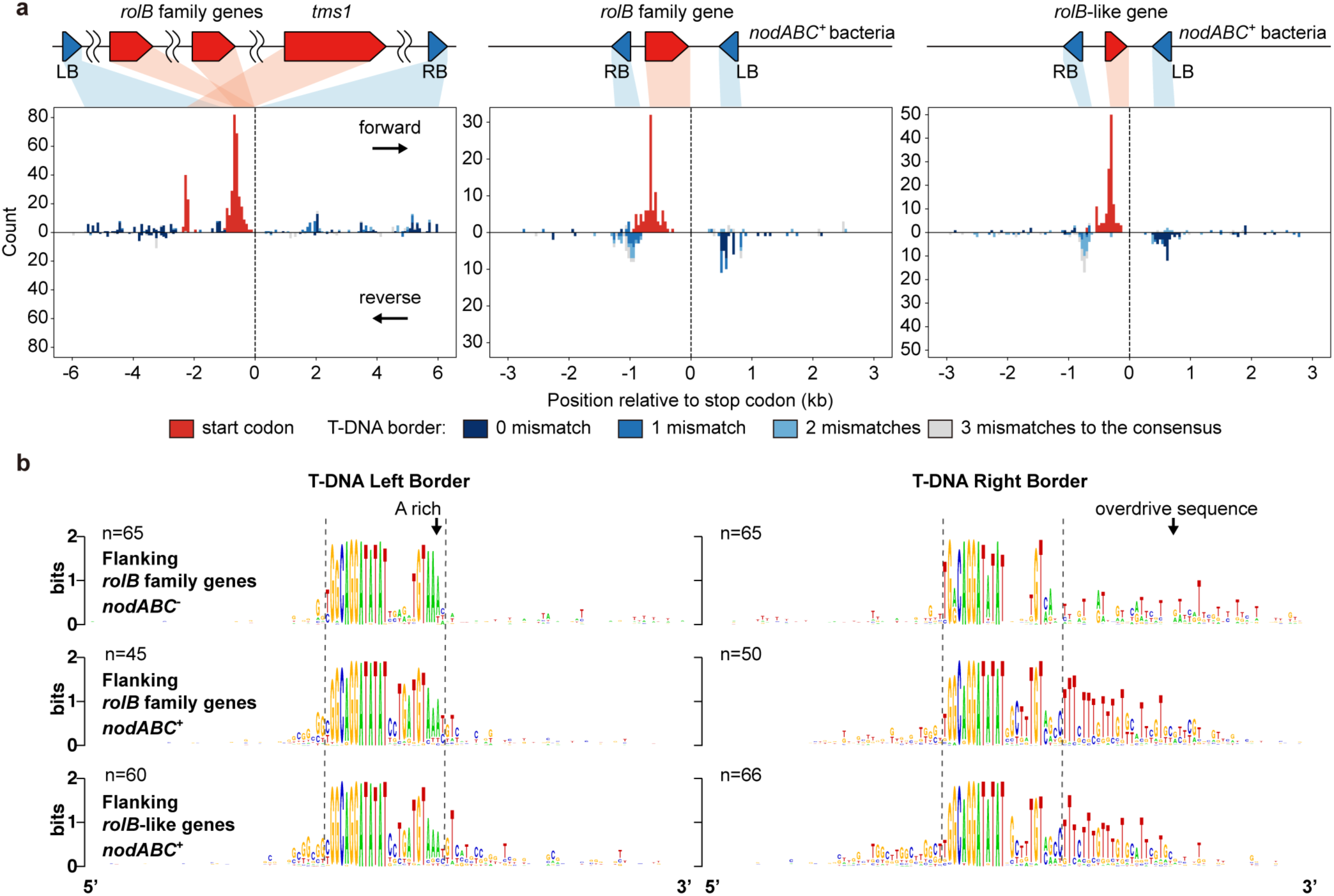
Positional organisation of start codons and T-DNA borders around *rolB/rolB*-like genes. **a**, Histograms showing the positions and orientations of start codons and adjacent T-DNA border-like sequences relative to the stop codons of non-redundant *rolB/rolB*-like genes. Left, *rolB* genes from genomes lacking the *nodABC* operon; middle, *rolB* genes from *nodABC*⁺ genomes; right, *rolB*-like genes. **b,** Consensus left and right T-DNA border sequences (25 bp), including 50 bp of upstream and downstream flanking sequence, for the three *rolB/rolB*-like gene groups shown in (**a**). The A-rich region of the left border and the T-rich ‘overdrive’ region downstream of the right border are indicated.

### Rhizobial mini-T-DNAs function as transferable, plant-expressed cargos in *Agrobacterium*

*Rhizobium etli* CFN42, a nitrogen-fixing symbiont of common bean (*Phaseolus vulgaris*), has been the only non-agrobacterial species shown to encode an endogenous, functional Vir-type transfer machinery, carried on its native p42a plasmid^25^. Earlier surveys detected no typical T-DNA cargos but identified five T-DNA border-like sequences, two of which flank an opine dehydrogenase gene and a trehalose-6-phosphate synthase gene^25,26^. Our analysis reveals two additional *rolB*-like genes within distinct mini-T-DNAs (Fig. 3a; T-DNAs are numbered according to their order in the direction of the adjacent *virB* operon).

**Fig. 3.**
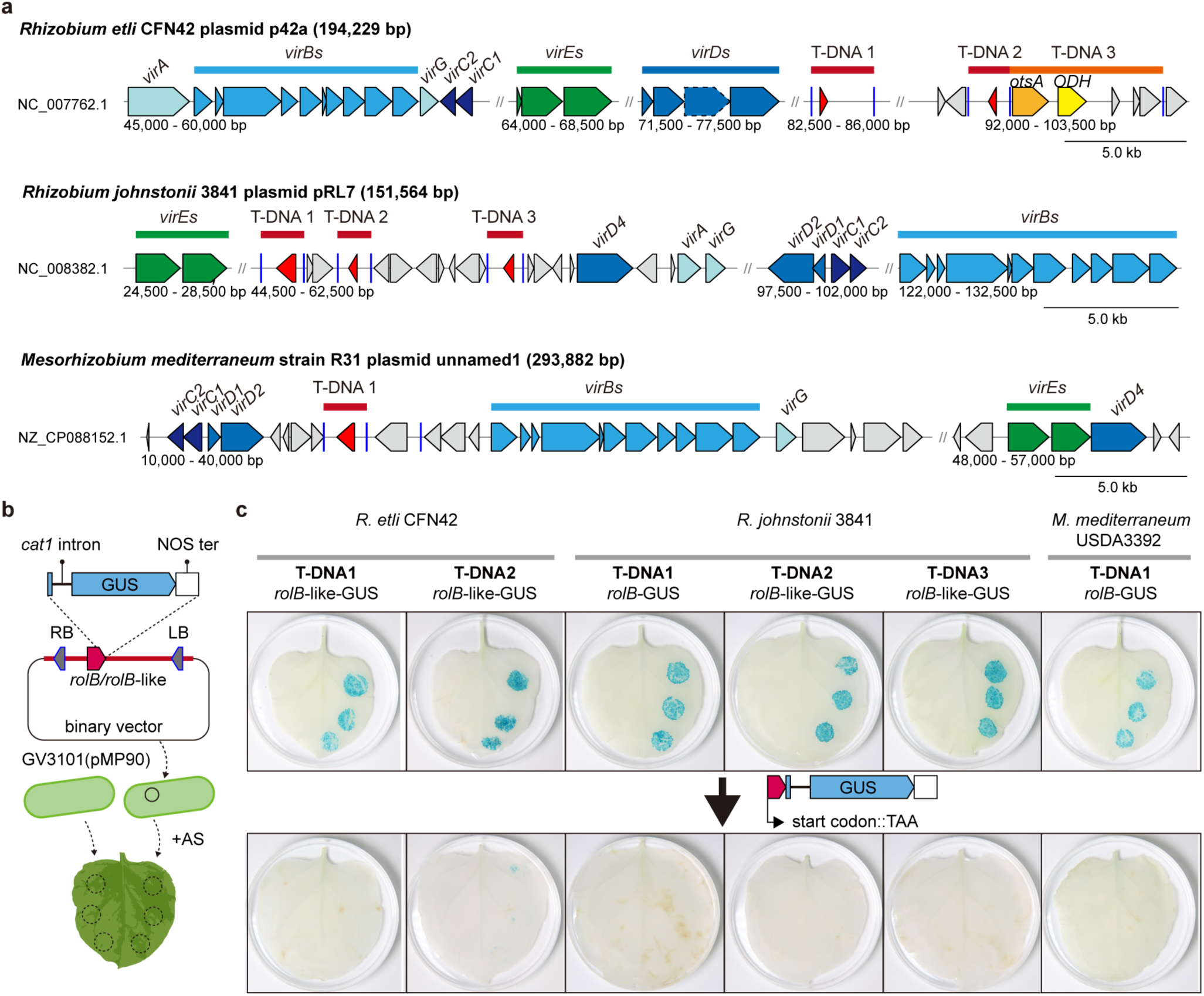
*Agrobacterium*-mediated transfer of rhizobial mini-T-DNAs drives expression of *rolB*/*rolB*-like genes in *Nicotiana benthamiana*. **a**, Genetic organisation of the mini-T-DNAs of three rhizobia, together with the homologues of *Agrobacterium* genes required for T-DNA transfer. All T-DNAs are plasmid-borne. (*Mesorhizobium mediterraneum* R31 is the sister strain of USDA3392 and has a complete genome assembly.) *otsA*, trehalose-6-phosphate synthase gene; *ODH*, opine dehydrogenase family protein gene. **b**, Schematic of the experimental setup. Each mini-T-DNA was inserted into a binary vector with the *rolB*/*rolB*-like gene translationally fused upstream of an intron-containing GUS reporter, and the constructs were introduced into *A. fabrum* GV3101(pMP90) for leaf-infiltration assays. AS, acetosyringone; LB, T-DNA left border; RB, T-DNA right border. **c**, The six T-DNAs containing *rolB*/*rolB*-like genes were transferable by *Agrobacterium* and drove expression of *rolB*/*rolB*-like genes in *N. benthamiana*. Mutation of the predicted start codons abolished GUS activity.

Given that *R. etli* CFN42 can recognize agrobacterial T-DNA, we reasoned that agrobacterial AMT machinery might also recognize rhizobial T-DNAs. We therefore selected mini-T-DNAs from three rhizobial strains: *R. etli* CFN42, *Rhizobium johnstonii* 3841, and *Mesorhizobium mediterraneum* USDA3392 for evaluation in *Agrobacterium* GV3101 (pMP90), a disarmed (T-DNA-deleted) derivative of *Agrobacterium fabrum* C58 that retains the *vir* transfer machinery. Although *R. johnstonii* 3841 was previously reported to lack detectable T-DNA transfer activity^25^, we identified three mini-T-DNAs in this strain (Fig. 3a).

To assess transfer and expression in plant cells, the six newly identified T-DNAs from these three strains were cloned into a binary vector, with an intron-containing GUS reporter translationally fused downstream of the *rolB/rolB*-like genes, and tested by agroinfiltration in *Nicotiana benthamiana* (*Nb*) leaves (Fig. 3b). All six constructs produced GUS signals in plant cells, indicating that the rhizobial T-DNAs were transferred and that their *rolB/rolB*-like cargos were expressed. Mutation of the predicted start codons abolished GUS signals, confirming translation of the fused ORFs. Notably, the *rolB*-like gene in T-DNA2 of *R. etli* CFN42 may utilize a non-canonical start codon (Extended Data Fig. 2a).

We further confirmed that the T-DNA borders and flanking regions of *R. etli* CFN42 were required to mediate transfer, as the GUS signal was detected upon addition of the border sequences from T-DNA1 and T-DNA2 (Extended Data Fig. 2b). Finally, we tested two genes located within the previously identified T-DNA3. The putative opine synthase gene was expressed in plants, whereas the trehalose-6-phosphate synthase gene showed no detectable expression (Extended Data Fig. 2c).

### Rhizobia mediate T-DNA transfer in root nodules

We next asked whether the three rhizobia themselves could mediate T-DNA transfer, given that most rhizobial T-DNAs identified here are located adjacent to *vir* genes. Using a betalain-producing RUBY reporter carried on an agrobacterial T-DNA^27^, we detected activity only for *R. etli* CFN42 on infiltration into *N. benthamiana* leaves (Extended Data Fig. 3a), consistent with a previous report^25^. We then introduced the same *rolB*/*rolB*-like-GUS translational fusion constructs into the three rhizobial strains and assayed T-DNA transfer under comparable conditions. Despite robust GUS expression when delivered by GV3101-mediated agroinfiltration (Fig. 3b), no GUS signal was detected for the three rhizobial strains following infiltration. Replacing the *rolB*-like-GUS fusion genes of *R. etli* CFN42 with a GUS gene under a strong CaMV 35S promoter likewise failed to produce detectable transformation (Extended Data Fig. 3).

Because *rolB*-like genes and a subset of *rolB* genes are largely restricted to rhizobia, which colonize host cells within root nodules, we hypothesized that T-DNA transfer might occur preferentially in this organ. To test this, we inoculated seedlings of the corresponding host plants with rhizobia carrying the *rolB/rolB*-like*-*GUS constructs, harvested root nodules after approximately three weeks, and assayed GUS activity. All three strains showed rhizobial T-DNA-dependent GUS expression in nodules, and the binary vector carrying the GUS gene under the 35S promoter did not produce the signal (Fig. 4). In all cases, signal was confined to nodules, with none detected in adjacent root tissue. These results demonstrate that T-DNA-carrying rhizobia can specifically transform the nodules of their native plant hosts.

**Fig. 4.**
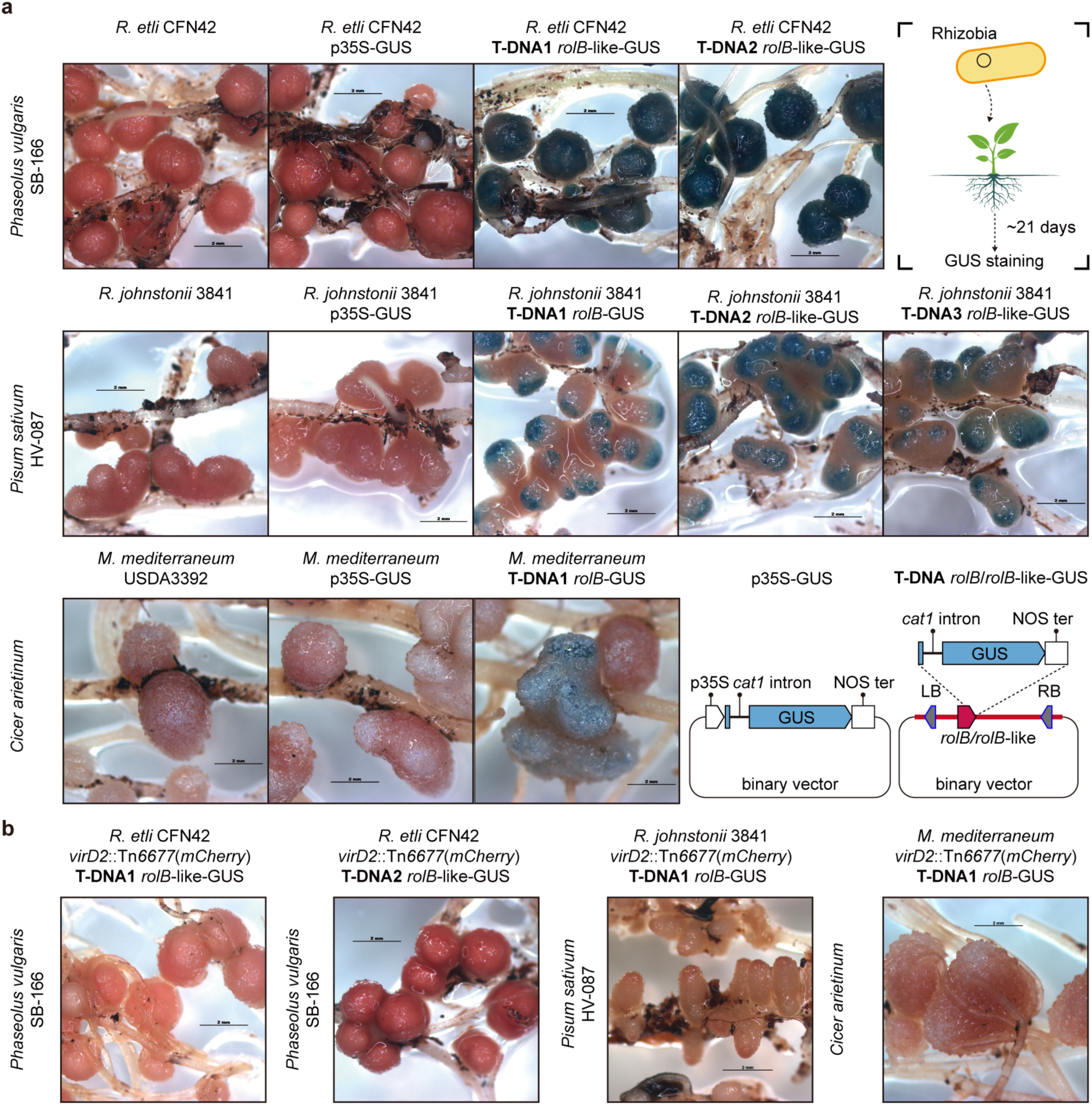
Rhizobia-mediated transfer of rhizobial mini-T-DNAs drives expression of *rolB*/*rolB*-like genes in root nodules of native host plants. **a**, GUS activity in root nodules formed by three rhizobial strains carrying binary vectors. Rows, top to bottom: *Rhizobium etli* CFN42 on common bean (*Phaseolus vulgaris* SB-166), *Rhizobium johnstonii* 3841 on pea (*Pisum sativum* HV-087), and *Mesorhizobium mediterraneum* USDA3392 on chickpea (*Cicer arietinum*). In each row, the first panel shows nodules formed by rhizobia lacking a binary vector and the second shows rhizobia carrying a binary vector with a p35S-GUS cassette but no T-DNA borders (negative controls); the remaining panels show nodules formed by rhizobia carrying that strain’s mini-T-DNAs, in which GUS is translationally fused downstream of the *rolB*/*rolB*-like gene, as in Fig. 3 and Extended Data Fig. 3. The cargo of each binary vector is indicated beneath the corresponding panel. Scale bars, 2 mm. Top right, schematic of the experimental setup; bottom right, schematic of the plasmids used. LB, T-DNA left border; RB, T-DNA right border; *cat1* intron, first intron of the castor bean (*Ricinus communis*) catalase gene; p35S, CaMV 35S promoter; NOS ter, nopaline synthase (NOS) terminator. **b**, The three rhizobial strains carrying a *virD2* disruption lost the ability to mediate T-DNA transfer into root nodules. Tn*6677*(*mCherry*), a CRISPR-guided transposon carrying a constitutive mCherry reporter. Scale bars, 2 mm.

Nodules formed by *R. etli* CFN42 on common bean and by *R. johnstonii* 3841 on pea (*Pisum sativum*) showed universal transformation, even without antibiotic selection to maintain the plasmids. By contrast, nodules formed by *M. mediterraneum* USDA3392 on chickpea (*Cicer arietinum*) showed sporadic GUS signal, detected in approximately one-third of nodules.

Notably, in the indeterminate nodules formed by *R. johnstonii* 3841 on pea, GUS activity was consistently restricted to the nodule tips across all three T-DNAs carrying *rolB/rolB*-like genes.

Disruption of *virD2* in all three strains abolished transformation, confirming its requirement for rhizobia-mediated transformation in nodules (Fig. 4c). Loss of T-DNA transfer did not impair nodulation, indicating that T-DNA delivery is dispensable for nodule formation under the conditions tested.

### Rhizobial RolB-like proteins perturb plant development

Having shown that rhizobial mini-T-DNAs are delivered into nodule cells and expressed, we asked whether the encoded RolB-like cargos are functional once inside plant cells.

Transgenic *Arabidopsis thaliana* overexpressing C-terminal mEGFP fusions of the RolB-like proteins from *R. etli* CFN42 (T-DNA2) and *R. johnstonii* 3841 (T-DNA3) with the 35S promoter displayed severe epinasty and stunted growth. This phenotype resembles that caused by the plast-family oncoprotein 6b^28,29^, consistent with perturbation of phytohormone balance (Extended Data Fig. 4). These results show that at least some rhizobial RolB-like cargos are biologically active *in planta*, supporting their role as functional T-DNA cargos.

### Rhizobial T-DNA transfer systems retain VirG-dependent activation but diverge in upstream signalling

Having examined the mini-T-DNAs, we next turned to how the transfer system itself is regulated. In agrobacteria, expression of the T-DNA transfer system is controlled by the VirA-VirG signalling pathway^30^. VirA is a transmembrane sensor kinase that detects plant-derived phenolic compounds and integrates additional cues, including monosaccharides and low pH, through interaction with ChvE^31–33^. Activated VirA phosphorylates the response regulator VirG, which then binds and activates *vir* gene promoters^34–36^. The T4SS and VirA-VirG system of *R*. *etli* CFN42 resemble those of agrobacteria^37^. By contrast, *R. johnstonii* 3841 and *M. mediterraneum* USDA3392 possess divergent signalling components, suggesting that distinct upstream pathways regulate their T-DNA transfer systems (Fig. 5a).

**Fig. 5.**
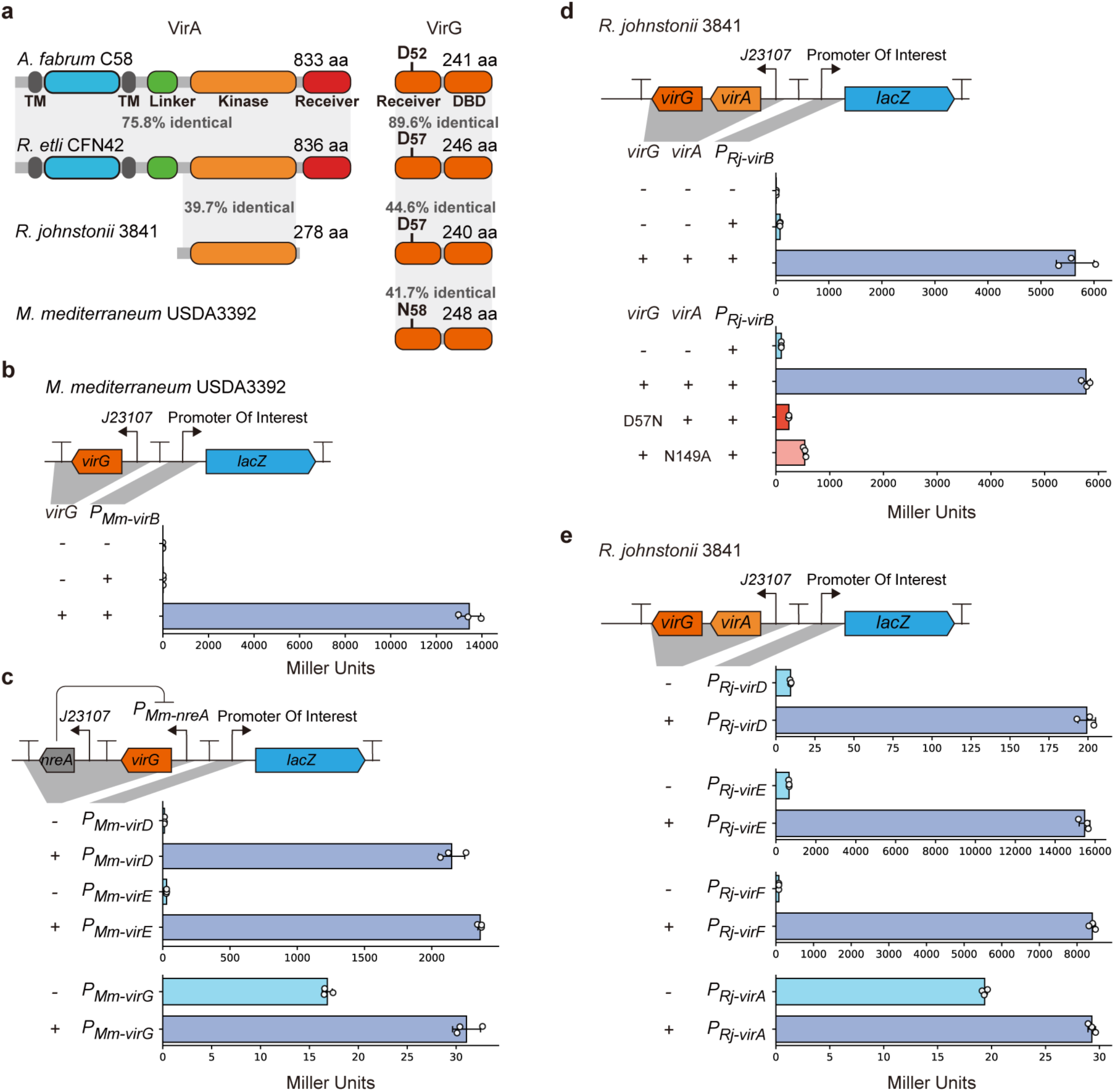
Rhizobial T-DNA transfer systems are under distinct transcriptional control. **a**, VirA and VirG homologues from the three rhizobial strains used in this study, compared with those of *Agrobacterium fabrum* C58. In *Rhizobium etli* CFN42, VirA and VirG have domain architectures highly similar to those of *A. fabrum*. In *Rhizobium johnstonii* 3841, VirA is truncated, retaining only the kinase domain. *Mesorhizobium mediterraneum* USDA3392 lacks VirA entirely, and the conserved phosphorylation site of its VirG is substituted by asparagine. **b**, Constitutive expression of VirG in *M. mediterraneum* USDA3392 activated the *virB* promoter. **c**, Leaky VirG expression in *M. mediterraneum* USDA3392 activated the *virD* and *virE* promoters, and more weakly the *virG* promoter itself. This basal expression caused a smaller growth defect than constitutive expression and was sufficient to activate the *virB* promoter (Extended Data Fig. 7). **d**, Constitutive expression of VirAG in *R. johnstonii* 3841 activated the *virB* promoter; mutation of either the VirG phosphorylation site or an active-site residue of VirA largely abolished this effect. **e**, Constitutive expression of VirAG in *R. johnstonii* 3841 also activated the *virD*, *virE*, and *virF* promoters. The *virA* promoter was weakly induced.

In *R. johnstonii* 3841, VirA is markedly shorter (278 aa), retaining only the kinase domain and part of the α-helical region predicted to mediate dimerization; a similar architecture is conserved among homologues with >60% amino acid identity. Disruption of either *virA* or *virG* abolished T-DNA transfer in nodules, confirming that both components are required for T-DNA transfer (Extended Data Fig. 5f). In *M. mediterraneum* USDA3392, *virA* is absent, and its VirG carries an asparagine substitution at the conserved phosphorylation-site aspartate; the same substitution is also present in close VirG homologues from other bacteria, suggesting that this group of regulators is unlikely to be activated by canonical phosphorylation (Fig. 5a).

To test whether these RMT systems respond to the signals that induce agrobacterial *vir* genes, we fused the *virB* promoter of each strain to *lacZ* and assayed its induction by the phenolic compound acetosyringone (AS) in *Agrobacterium*-inducing buffer (AIB), a glucose-containing buffer at low pH. In contrast to agrobacterial GV3101 *virB* promoter that is highly induced by acetosyringone, none of the three *virB* promoters was induced by acetosyringone (Extended Data Fig. 5a). For *R. johnstonii* 3841 and *M. mediterraneum* USDA3392, this is consistent with their divergent regulators. For *R. etli* CFN42, whose regulators closely resemble those of agrobacteria, the result was more unexpected, given earlier work showing that its *virE* promoter is mildly induced by acetosyringone^37^. Instead, the CFN42 *virB* promoter was already highly active in AIB in the absence of acetosyringone (Extended Data Fig. 5a), while remaining at background levels in TY medium (Extended Data Fig. 5b). To identify the relevant signal, we assayed TY medium buffered across a range of pH values, with or without glucose, and found that low pH and glucose alone were sufficient for activation (Extended Data Fig. 5c). This differs from agrobacteria, in which low pH and specific monosaccharides enhance *vir* promoter activity but cannot substitute for a phenolic inducer. The *R. etli* CFN42 *virB* promoter was also activated by arabinose but not fructose, a specificity reminiscent of the ChvE-dependent sugar enhancement of VirA signalling in agrobacteria (Extended Data Fig. 5d). At low pH in TY medium, glucose also activated the *virD*, *virE*, and *virA* promoters (Extended Data Fig. 5e), with the strongest response at the *virD* operon and only modest induction of *virA*. (Extended Data Fig. 5e).

Strains *R. johnstonii* 3841 and *M. mediterraneum* USDA3392, with more divergent regulators, their *virB* promoters were expressed at low levels and did not respond to acetosyringone and we could not identify an inducing signal for them. Thus, we asked instead whether their divergent regulators are themselves sufficient to activate core *vir* gene expression. We assayed transcriptional fusions of their *vir* promoters to *lacZ*, with and without co-expression of the corresponding regulators. In *M. mediterraneum* USDA3392, which lacks VirA, constitutive expression of VirG from J23107 (a synthetic constitutive promoter; BBa_J23107, iGEM Registry of Standard Biological Parts) strongly activated the *virB* promoter (Fig. 5b) but caused a growth defect. To reduce this toxicity, we placed *virG* under the Ni²⁺-responsive NreA regulatory system, in which NreA represses its own promoter and dissociates upon Ni²⁺ binding^38^. Basal expression from this system, even without Ni²⁺ induction, was sufficient to activate the *virB* promoter while substantially reducing the growth defect (Extended Data Fig. 6a), and was therefore used for subsequent experiments. VirG-dependent induction was also observed for the *virD* and *virE* promoters, and, weaklier, the *virG* promoter itself (Fig. 5c).

Similarly, constitutive expression of the VirA-VirG pair from *R. johnstonii* 3841 strongly activated the *virB* promoter. Mutation of either a putative catalytic residue in VirA or the phosphorylation site in VirG largely abolished activation, indicating that this truncated two-component system remains phosphorylation-dependent (Fig. 5d). Induction was also observed for the *virD*, *virE*, and *virF* promoters, and, weaklier, the *virA* promoter itself (Fig. 5e). Because these assays were performed in strains carrying intact endogenous *virA-virG* or *virG* loci, these results suggest that endogenous regulator expression is low under the tested conditions. Additionally, the divergent rhizobial regulators could activate rhizobial *virB* promoters in the GV3101 background independently of their native cellular context. The result supported the sufficiency of these regulators for *vir* promoter activation (Extended Data Fig. 6b-c).

We next asked whether the *vir* genes responsible for rhizobial T-DNA transfer systems respond to symbiotic flavonoid signalling. In the *M. japonicum* R7A effector-delivery system, whose VirB protein is 80% identical to those of agrobacteria, *virA* expression is controlled by NodD and induced by the flavonoid naringenin^39^, which subsequently activates *virB* and effector genes. Because naringenin also induces nodulation genes in *R. johnstonii* 3841 and many other rhizobia^40–42^, we tested whether naringenin activates the *virA*, *virG*, or *virB* promoters of the three rhizbobial strains. As a positive control confirming that the treatment was effective, the *nodA*/*nodABC* promoters of all three strains were strongly induced by naringenin; none of the *vir* promoters showed detectable induction (Extended Data Fig. 6d-f). These systems are therefore not directly controlled by canonical NodD-dependent flavonoid signalling under the conditions tested.

Together, these findings show that the divergent rhizobial VirA-VirG and VirG-only systems are nonetheless sufficient to activate core *vir* gene expression, yet their upstream inputs differ from those of agrobacteria. The *virB* promoters of *R. johnstonii* 3841 and *M. mediterraneum* USDA3392 were induced neither by the acetosyringone that triggers the agrobacterial *vir* regulon nor by NodD-dependent flavonoid signalling. In *R. etli* CFN42, whose VirA-VirG most closely resembles that of agrobacteria, the *virB* promoter was activated by the acidic pH and specific monosaccharides (e.g. arabinose and glucose), cues that are enriched in planta. Rhizobial T-DNA transfer systems therefore appear to retain conserved, VirG-dependent activation of the core transfer genes while being wired to distinct upstream signals.

### Rhizobial T-DNA transfer systems retain ancestral-like *virB* operon architectures

Although the VirB proteins of *R. etli* CFN42 closely resemble their agrobacterial counterparts, those of *R. johnstonii* 3841 and *M. mediterraneum* USDA3392 are markedly divergent^25^. Their VirB4 proteins share <30% amino acid identity with agrobacterial VirB4, even lower than the ∼40% identity observed in the *Bartonella henselae* T4SS effector-delivery system. This divergence is unexpected in functional T-DNA transfer systems, particularly since the VirD2 relaxase domains from all three rhizobia remain highly conserved, sharing nearly 60% identity with *A. fabrum* C58.

These divergent rhizobial systems also exhibit distinct *virB* operon architectures. We identified putative *virB7* in *R. johnstonii* 3841 and *M. mediterraneum* USDA3392, located between *virB4* and *virB5*, rather than at the canonical position between *virB6* and *virB8*. PhoA assays in *Escherichia coli* confirmed that these rhizobial *virB7* genes encode functional export signals (Extended Data Fig. 7)^43^. Despite these differences in homology and gene order, promoter assays and nodulation experiments support their role in T-DNA transfer: *virB* genes were co-induced with *virD* and *virE*, and disruption of *virB4* in *R. johnstonii* 3841 abolished T-DNA transfer in nodules (Extended Data Fig. 5f).

To investigate the evolutionary basis of these differences, we analysed alphaproteobacterial protein-secretion type IV secretion systems (pT4SSt; the VirB/VirD4 lineage; MPF type T) identified using TXSScan (MacSyFinder)^44,45^. For phylogenetic analysis, we clustered the VirB4 sequences of these systems at 80% amino acid identity and used the resulting representatives. The analysis resolved two distantly related clades, whose loci co-localise with *rolB* or *rolB*-like genes (Extended Data Fig. 8a): clade A, which includes most agrobacterial T-DNA transfer systems and the *R. etli* CFN42 system; and clade B, which includes the divergent rhizobial systems of *R. johnstonii* 3841 and *M. mediterraneum* USDA3392.

Resolving clades A and B at higher resolution (95% amino acid identity clustering) suggested that agrobacterial *virB* operon was acquired through replacement of the entire operon, whereas many rhizobial systems retain a distinct, ancestral-like *virB* operon in gene content and arrangement (Fig. 6). Among clade A systems, *virD2* co-localised with *virB4* largely in canonical AMT systems and their close relatives. Basal clade A lineages not associated with T-DNA transfer instead co-localised with MobF-family relaxase genes rather than *virD2*. Clade B *virB4* frequently co-localised with *virD2*, including in distantly related lineages. These patterns suggest that the association between *virD2* and clade B *virB4* predates the diversification of many rhizobial T-DNA transfer systems.

**Fig. 6.**
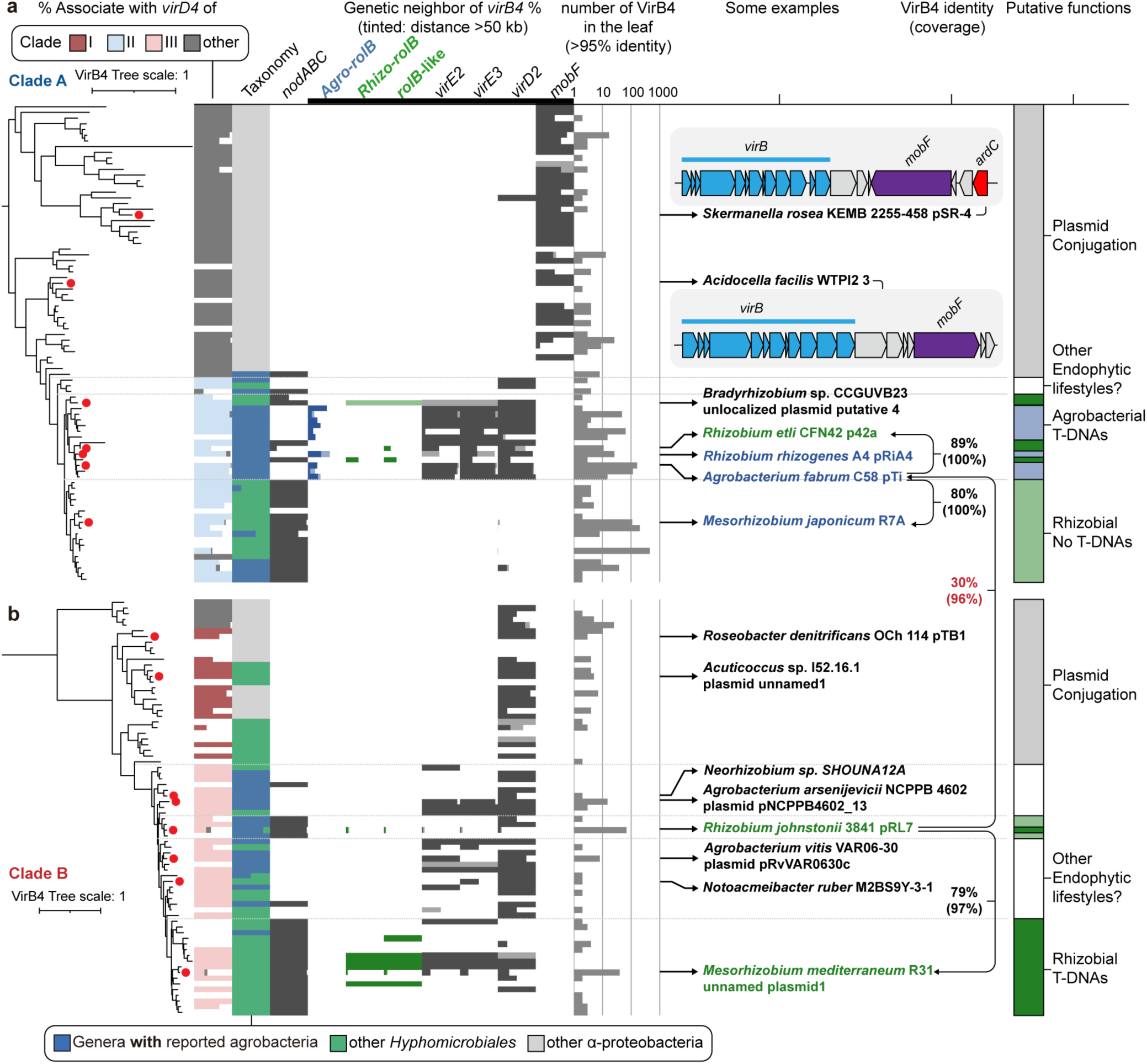
Phylogenetic analyses of alphaproteobacterial VirB4 proteins associated with agrobacterial and rhizobial T-DNA transfer systems. **a, b**, Phylogenetic trees of alphaproteobacterial VirB4 proteins, constructed from representative sequences of clusters sharing >95% amino acid identity. For each cluster, co-localisation of *virB4* with clade I, II, or III *virD4* loci, as defined in Extended Data Fig. 8, is indicated by colour, and genus-level taxonomic composition is colour-coded according to NCBI taxonomy. The proportion of clusters whose genomes contain a *nodABC* gene cluster is shown in dark grey. The presence of selected genes on the same contig as *virB4* is indicated similarly, with tinted colours denoting genes located >50 kb from *virB4*. *rolB* genes are separated into two groups corresponding to the clades primarily associated with rhizobia or with agrobacteria, as defined in Fig. 1c. Cluster sizes are shown as bar charts on a logarithmic scale. VirB4 proteins are indicated for the rhizobial strains analysed in this study and for the well-characterized systems of *Agrobacterium fabrum* C58 (pTiC58), *Rhizobium rhizogenes* A4 (pRiA4), and *Mesorhizobium japonicum* R7A; selected VirB4 proteins from putative endophytic T-DNA transfer systems that are neither oncogenic nor rhizobial are also labelled. Pairwise identities between selected VirB4 proteins are shown as connecting lines, and putative functions of the T4SS subclades are indicated at right. **a**, VirB4 phylogeny of the clade closely related to classical Ti-plasmid T-DNA transfer systems (clade A in Extended Data Fig. 8a), rooted as in Extended Data Fig. 8a. *ardC*, anti-restriction protein ArdC, commonly associated with plasmid-conjugation genes. **b**, VirB4 phylogeny of the clade distantly related to classical Ti-plasmid T-DNA transfer systems (clade B in Extended Data Fig. 8a), rooted as in Extended Data Fig. 8a.

The VirD4 protein phylogeny provided additional support for this model. Transfer-associated VirD4 proteins formed a single clade of two subclades, II and III, whose closest non-transfer sister branch, clade I, predominantly co-localised with clade B *virB4* loci (Extended Data Fig. 9). Within the transfer-associated clade, clade II co-localised with clade A *virB4* loci and clade III with clade B *virB4* loci. Features of clade I systems, including nearby T-DNA border-like sequences that may act as *oriT* sites and anti-restriction genes near the *virD* loci, are consistent with an ancestral role in plasmid conjugation and imply an early association with clade B *virB4*.

This concordant phylogenetic and co-localisation pattern is consistent with a *virB* replacement event early in the evolution of T-DNA transfer systems, giving rise to VirD4 clade II, which includes canonical AMT systems and *M. japonicum* R7A-like effector-delivery systems. In contrast, many RMT systems contain VirD4 clade III and retain clade B *virB* modules. Together, the VirB4 and VirD4 phylogenies support a modular-replacement model in which plasmid-borne VirB4 clade B-associated systems resemble an ancestral-like configuration, whereas clade A modules were recruited into canonical AMT and related systems. The analysis further revealed many additional putative T-DNA transfer systems, recognizable from their association with *virB*, *virD*, and *virE* homologues and candidate T-DNA regions (examples in Extended Data Fig. 8b). This suggests that AMT and RMT represent only two subsets of a much broader diversity of such systems among plant-associated bacteria.

## Discussion

T-DNA transfer has historically been viewed through the lens of agrobacterial virulence, with AMT treated as a unique evolutionary innovation for pathogenesis. Here we show that this view is incomplete. Homologous T-DNA transfer systems occur well beyond canonical oncogenic plasmids associated with pathogenic agrobacteria, and are found in diverse nitrogen-fixing rhizobia. These newly discovered RMT systems were experimentally validated in three rhizobia-host pairs as functional for nodule-specific transformation. These and other findings, including a genome-wide T-DNA screen, the discovery of rhizobial *rolB*/*rolB*-like genes, and phylogenetic analysis of the transfer machinery, recast bacteria-to-plant DNA transfer as an older and more widespread capability, of which agrobacterial pathogenesis and rhizobial symbiosis represent two specialized outcomes.

Agrobacteria do not form a monophyletic group: the tumorigenic lifestyle has arisen repeatedly in lineages that are often more closely related to non-pathogenic rhizobia than to one another^18^. Given this, and the simplicity of the rhizobial mini-T-DNAs relative to the larger, multi-gene T-DNAs of agrobacteria, it is tempting to regard RMT systems as ancestral to AMT. The picture, however, is more complex. Two lines of evidence pull in opposite directions. The rhizobial RolB-like proteins occupy a basal position within the RolB/RolB-like proteins, consistent with an ancestral-like RMT. Yet the canonical rhizobial RolB proteins are less diverse than their agrobacterial counterparts, the opposite of what an older, longer-diversifying rhizobial lineage would predict, although diversity reflects selection, lineage expansion, sampling, and later horizontal gene transfer as much as age, so the contrast is indicative rather than conclusive. Although AMT systems arose from modular replacement of *virB* operon, retention of an ancestral-like *virB* configuration by the majority of RMT systems is not by itself sufficient to establish RMT as the AMT ancestor. It also remains unclear why all known AMT systems appear to derive from a single *virB* replacement event; whether this reflects incomplete sampling of AMT diversity or selective constraints requires further study.

Phylogenetic analysis revealed that RMT systems are distributed across the VirB4 tree, with some carrying AMT-like *virB* genes. We therefore cannot rule out that the current association of RMT systems with an ancestral-like *virB* configuration reflects later horizontal transfer. Furthermore, the VirAG/VirG proteins of the three rhizobial strains we tested are more closely related to homologues in non-RMT systems than to one another. Together, these observations indicate that, like AMT systems, the different modules of RMT systems have also been shaped by recombination-driven mosaicism^19^. The same process is evident in the type V pTi, in which pRi-like *vir* genes are associated with crown gall-inducing T-DNAs^18^. We cannot exclude that an early RMT-like system, bearing mini-T-DNAs with *rolB*/*rolB*-like genes, predated and gave rise to the first oncogenic AMT system. But taken together, these signals suggest that the extant RMT and AMT systems are better understood as co-descendants of a shared pool of ancestral systems, shaped by ongoing HGT and selection, than as one having given rise to the other.

Several questions remain open. Only a minority of rhizobia carry RMT systems, and in the three strains tested here RMT was dispensable for nodulation under laboratory conditions, leaving its role in the symbiosis, the functions of the rhizobial RolB/RolB-like cargos, and the signals that trigger transfer to be defined. Identifying those signals will be central to understanding when transfer occurs during symbiosis and how a conserved secretion system becomes integrated into different plant-associated lifestyles.

Beyond these questions, RMT offers a tool for studying nodulation and a possible route for plant engineering with properties distinct from AMT. Legumes are often recalcitrant to AMT, whereas RMT operates natively in these hosts. Many rhizobia also nodulate only compatible legume species, a specificity that could provide a layer of biological containment unavailable with the broad host range of agrobacteria. Because transgene expression was detected only in nodules, RMT may allow new traits to be directed to the tissue responsible for nitrogen fixation. Whether T-DNA transfer is confined to nodule cells remains to be tested, but if it is, other organs including seeds would remain unmodified, which would bear directly on how such plants are regulated.

Consistent with the possibility that RMT is already active in the field, we detected RMT systems in commercial legume inoculants, including *Ensifer meliloti* RRI128 and *M. mediterraneum* R31, suggesting that such transfer may already occur in agricultural symbioses. Realizing these applications will require defining transfer efficiency, host range, and tissue specificity directly, but the distinct properties of RMT make it a promising complement to established transformation systems.

More broadly, our findings show that bacteria-to-plant DNA transfer is not restricted to pathogenic agrobacteria, establish RMT as a symbiosis-associated counterpart of AMT, and point to a wider evolutionary diversity of T-DNA transfer systems among plant-associated bacteria.

## Methods

### Identification of candidate T-DNAs in alphaproteobacterial genomes

Alphaproteobacterial genomes were downloaded from NCBI on 10 December 2025. In total, 63,759 genomes were retrieved, of which 38,231 were annotated. Putative T-DNA borders were identified using the degenerate consensus sequence YGRCAGGATATATNNNNNKGTMAWN. Hits with fewer than four mismatches were retained. Sequences shorter than 50 kb flanked by T-DNA borders in the same orientation were considered candidate T-DNAs.

Because close homologues of *virD2* are rarely found outside the order Hyphomicrobiales, candidates identified outside Hyphomicrobiales were used as negative controls to define bona fide T-DNAs. Candidate regions with a coding sequence density below 80% and a GC content at least 7% lower than that of the host genome were classified as likely T-DNAs.

### Identification of virulence-related proteins and type IV secretion systems

Proteins of interest were identified using hmmsearch with the following Pfam profiles: PF02027 (RolB family proteins), PF07229 (VirE2), PF06661 (VirE3), PF03432 (VirD2), and NF041492.1 (MobF). RolB-like proteins were identified using a custom HMM profile constructed from homologues of WP_254106469.1. The distant homology between WP_254106469.1 and RolB family proteins was detected using HHpred^46^. Homologues were retrieved using PSI-BLAST, which reached convergence after five iterations. The alignment of 44 sequences was used to build the HMM profile using hmmbuild, after removing two truncated sequences.

RolB and RolB-like hits with a bit score greater than 25, or greater than 20 when located within predicted T-DNAs, were considered bona fide hits. For all other proteins, a bit score cutoff of 25 was used. In addition, for RolB and RolB-like proteins, six-frame translations of unannotated genomes generated using SeqKit2^47^ were screened.

Protein-secretion type IV secretion systems (pT4SSt) were identified using MacSyFinder with the TXSScan pT4SSt model^44,45^. pT4SSt systems co-localised with *rolB* or *rolB*-like genes were considered candidate *virB* loci.

### Structural prediction and analysis of RolB and RolB-like proteins

Protein structures of RolB and RolB-like proteins were predicted using AlphaFold^48^. Predicted structures were compared and visualized in ChimeraX^49^. Topology diagrams were generated using PDBsum^50^.

### Identification of *nodABC* clusters

The presence of the *nodABC* gene cluster was used as an approximate indicator of rhizobia. Genes encoding NodA, NodB, and NodC were identified using hmmsearch with the HMM profiles PF02474, TIGR04243, and TIGR04242. A *nodABC* cluster was defined when *nodA*, *nodB*, and *nodC* were present within a 30 kb genomic region. A 30 kb window was used because, although *nodABC* commonly forms an operon, in some bacteria (e.g., *Rhizobium etli* CFN42 and many *Mesorhizobium*) *nodA* is located distant from *nodB* and *nodC*.

### Phylogenetic analysis of Alphaproteobacterial mating pair formation type T VirB4/VirD4 proteins

For making VirB4 trees, VirB4 protein sequences were extracted from MacSyFinder^44^ hits and clustered using MMseqs2 (v18.8cc5c)^51^ with an 80% sequence identity cutoff.

Representative sequences from each cluster were aligned using MUSCLE (v5.1)^52^, and a phylogenetic tree was constructed with FastTree (v2.1.10, SSE3)^53^ under the WAG (Whelan and Goldman) substitution model. Two clades associated with *rolB* and *rolB*-like genes were subsequently extracted and re-clustered using MMseqs2 at a 95% identity cutoff. Phylogenetic trees were then reconstructed following the same procedure. Trees were visualized using the Interactive Tree Of Life (iTOL) v7.5^54^. VirD4 phylogenetic analysis was performed similarly.

Sequences were first clustered at 80% identity and used to construct a tree from representative sequences. The clade containing agrobacterial VirD4 and its sister lineages was then extracted, re-clustered at 95% identity, and used to construct the tree.

### Bacterial strains and culture conditions

Bacterial strains and plants used in this study are listed in Supplementary Table 1. *Rhizobium etli* CFN42 and *Rhizobium johnstonii* 3841 were obtained from the Leibniz Institute DSMZ–German Collection of Microorganisms and Cell Cultures (Braunschweig, Germany), and *Mesorhizobium mediterraneum* USDA3392 from the Bioresource Collection and Research Center (BCRC, Hsinchu, Taiwan). Rhizobia were cultured in TY medium (0.5% (w/v) tryptone, 0.3% (w/v) yeast extract and 10 mM CaCl₂) at 28 °C. *Agrobacterium* GV3101 (pMP90) was cultured in LB medium (1% (w/v) tryptone, 0.5% (w/v) yeast extract and 0.5% (w/v) NaCl) at 28 °C, and *Escherichia coli* DH5α and S17-1 in LB medium at 37 °C. Solid media were prepared with 1.5% (w/v) agar. Liquid cultures were grown in 5 ml medium in 15 ml capped tubes at 200 r.p.m. Where required, media were supplemented with kanamycin (50 μg ml⁻¹), spectinomycin (50 μg ml⁻¹), chloramphenicol (25 μg ml⁻¹), streptomycin (500 μg ml⁻¹) or nalidixic acid (20 μg ml⁻¹).

### Plasmid construction and rhizobia transformation

The plasmids used in this study are listed in Supplementary Table 2. DNA fragments were amplified using PrimeSTAR Max DNA Polymerase Ver. 2 (Takara). Bacterial genomic DNA templates were extracted using the Wizard® Genomic DNA Purification Kit (Promega). Plasmids were constructed by standard restriction cloning or using the NEBuilder HiFi DNA Assembly Master Mix (New England Biolabs). Colony PCR was conducted using the AppEx Taq DNA Polymerase.

*R. etli* CFN42 was transformed by electroporation (2.5 kV, 0.2 cm gap cuvette, single pulse; program Ec2) using a MicroPulser electroporator (Bio-Rad), recovered in TY medium for 4-6 hours at 28 °C, and transformants were selected on TY plates containing kanamycin or spectinomycin. *R. johnstonii* 3841 and *M. mediterraneum* USDA3392 were transformed by biparental conjugation with *E. coli* S17-1 as donor at a donor-to-recipient ratio of 1:1, mated on TY plates overnight at 28 °C. Transconjugants were selected on TY plates containing kanamycin and streptomycin (*R. johnstonii* 3841) or chloramphenicol and nalidixic acid (*M. mediterraneum* USDA3392); streptomycin and nalidixic acid counterselected against the donor.

### Bacterial gene disruption

Genes of interest were disrupted using the CRISPR-guided transposon-based INTEGRATE system. A derivative carrying the pVS1 replicon, pEA186, was used for delivery. pEA186 was a gift from Kan Wang (Addgene plasmid #187874; RRID: Addgene_187874)^55^.

CRISPR spacers targeting the genes of interest were designed using the CAST guide RNA tool^56^ and cloned into the CRISPR array of pEA186. The resulting plasmids were introduced into bacteria by transformation. Single colonies were screened for transposon insertion using colony PCR. Positive colonies were re-streaked and re-screened to confirm insertion. Colonies that showed the insertion signal but lacked the wild-type allele were selected. To cure the plasmid, bacteria were then cultured in antibiotic-free medium and plated on solid medium supplemented with 5% (w/v) sucrose.

### Leaf infiltration

The method is modified from the standard protocol^57^. GV3101 carrying the indicated plasmids was cultured in YEP medium (1% w/v yeast extract, 1% w/v bactopeptone, and 0.5% w/v NaCl) supplemented with appropriate antibiotics for 1-2 days at 28 °C with shaking at 200 r.p.m. until late log phase. Then the bacteria were diluted 50-fold into YEP medium supplemented with 10 mM MES and grown until the OD₆₀₀ reached approximately 1. Cells were harvested by centrifugation and resuspended in MMA medium (2% w/v sucrose, 0.5% w/v MS salts, 1.95 g l⁻¹ MES, and 20 µM acetosyringone, pH 5.6) to an OD₆₀₀ of 0.3. After incubation at room temperature (∼25 °C) for 1-6 h, the bacterial suspension was infiltrated into the abaxial side of 3-week-old *Nicotiana benthamiana* leaves. Leaves were harvested for analysis 3-6 days after infiltration.

### Nodulation assays

Legume seeds were surface-sterilized with 75% ethanol for 10 min, followed by 95% ethanol for 10 min. Sterilized seeds were then soaked in sterile Milli-Q water for 1 h until swollen and subsequently planted in sterilized soil. Common bean and chickpea plants were grown at 25 °C under a 16 h light/8 h dark photoperiod, whereas pea plants were grown at 22 °C under the same photoperiod.

When plants developed the first pair of true leaves, they were inoculated with rhizobia. Rhizobial cultures were grown in liquid medium with appropriate antibiotics to late log phase, harvested by centrifugation, washed with Milli-Q water, and diluted to an OD₆₀₀ of 0.1 in Milli-Q water. Each plant was inoculated with 10 ml of the bacterial suspension. Root nodules were harvested for analysis approximately 3 weeks after inoculation.

### GUS and Gal staining of leaves and root nodules

For GUS staining, *Nicotiana benthamiana* leaves were immersed in GUS staining buffer containing 50 mM sodium phosphate buffer (pH 7.0), 2 mM K₃Fe(CN)₆, 2 mM K₄Fe(CN)₆, 0.1% Triton X-100, and 1 mM X-Gluc (5-bromo-4-chloro-3-indolyl-β-D-glucuronide). Leaves were vacuum-infiltrated until they became semi-transparent and then incubated in the dark at 37 °C for 4-8 h with gentle agitation at 60 r.p.m. After staining, leaves were washed once with 70% ethanol and then repeatedly with 95% ethanol under agitation until chlorophyll was removed. Root nodules were stained using the same procedure, except that ethanol clearing was omitted.

For Magenta-Gal staining following GUS staining, root nodules were first washed with 50 mM sodium phosphate buffer (pH 7.0) and incubated in the same buffer at 70 °C for 1 h.

Nodules were then vacuum-infiltrated, immersed in Magenta-Gal staining buffer containing 50 mM sodium phosphate buffer (pH 7.0), 2 mM K₃Fe(CN)₆, 2 mM K₄Fe(CN)₆, 0.1% Triton X-100, and 1 mM Magenta-Gal (5-bromo-6-chloro-3-indolyl-β-D-galactopyranoside), and incubated in the dark at 65 °C for 6-24 h with gentle agitation at 60 r.p.m.

### β-galactosidase promoter assay

Reporter plasmids carrying *lacZ* under the control of the promoter of interest, with or without constitutive co-expression of *virA*-*virG* (or *virG* alone, as indicated), were introduced into the relevant bacterial strains. Transformants were grown in the indicated medium supplemented with the appropriate antibiotics. At late logarithmic phase, 200 μl of culture was subcultured into 10 ml of fresh medium containing antibiotics and, where indicated, inducers, and incubated for 12–72 h (strain-specific times below). Promoter activity was then measured by a standard Miller assay and expressed in Miller units.

For assays of constitutive *virA*-*virG* expression, conditions were: *A. fabrum* GV3101(pMP90), YPD medium, ∼12 h; *R. johnstonii* 3841, TY medium, ∼12 h; *M. mediterraneum* USDA3392, TY medium, ∼48 h. For assays of medium composition and inducers, conditions were: *A. fabrum* GV3101(pMP90), *Agrobacterium*-inducing buffer (AIB), ∼24 h; *R. etli* CFN42, AIB or TY medium, ∼24 h; *R. johnstonii* 3841, AIB or TY medium, ∼24 h; *M. mediterraneum* USDA3392, AIB, ∼72 h, or TY medium, ∼48 h. Where TY medium was buffered to defined pH, 50 mM MES was used for pH 5.6 and 6.7 and 0.1 M Tris-HCl for pH 7.4. AIB consisted of 1xAB salts (18.7 mM NH_4_Cl, 1.2 mM MgSO_4_, 2.0 mM KCl, 0.09 mM CaCl_2_, and 0.009 mM FeSO_4_), 2 mM KH_2_PO_4_, 50 mM MES, 0.5% (w/v) glucose, and adjusted to pH 5.6 with KOH^58^. Inducers were added to the following final concentrations: naringenin, 10 μM; acetosyringone, 100 μM; glucose, arabinose, or fructose, each 0.5% (w/v).

### PhoA assay of putative VirB7

The *phoA* gene of *E. coli*, lacking its native signal peptide (*phoA′*), was amplified from pRMCD28 and assembled into the pBBR1MCS-2 backbone under the control of the *lacZ* promoter. A HindIII restriction site was introduced at the 5′ end of *phoA′*. pRMCD28 was a gift from Renato Morona (Addgene plasmid #167556; http://n2t.net/addgene:167556; RRID:Addgene_167556)^59^. Candidate *virB7* genes were cloned upstream of *phoA′*. PhoA activity was assessed by streaking *E. coli* strains carrying the plasmids onto LB agar plates supplemented with 40 µg ml⁻¹ 5-bromo-4-chloro-3-indolyl phosphate (XP) and 50 µg ml⁻¹ kanamycin.

### Generation and screening of transgenic *Arabidopsis* expressing rhizobial RolB/RolB-like-mEGFP fusions

The *rolB* and *rolB*-like genes were fused in frame at their C termini to mEGFP and cloned downstream of the CaMV 35S promoter in pCAMBIA1201, into which a kanamycin-resistance marker had been inserted for selection in bacteria. The resulting binary vectors were introduced into *A. fabrum* GV3101(pMP90), which was then used to transform *Arabidopsis thaliana* Columbia-0 by the standard floral-dip method. Primary transformants (T1) and their progeny (T2) were selected on hygromycin-containing medium. For each construct, T3 plants from three independent lines were grown on soil (16 h light/8 h dark, 22 °C) alongside wild-type controls for approximately three weeks, then assessed for GFP fluorescence and photographed to document growth phenotypes.

## Author Statements Acknowledgements

We thank the Cell Biology Core and the Genomic Technology Core at the Institute of Plant and Microbial Biology (Academia Sinica, Taiwan) for technical assistance.

## Author contributions

SCH and CHK conceptualized the project and drafted the manuscript. SCH designed and performed all experiments, analyses, and data visualization. EML contributed intellectual discussion and experimental designs. CHK acquired funding and supervised the project. All authors edited the manuscript and approved the submitted version.

## Usage of artificial intelligence tools

The authors used ChatGPT-5 and Claude Opus 5 to support brainstorming of manuscript organization, to suggest wording improvements, and to correct grammar. All substantive writing and the final text are the authors’ own work. No generative artificial intelligence tools were used in the preparation of figures or tables.

## Extended data legends

**Extended Data Fig. 1.**
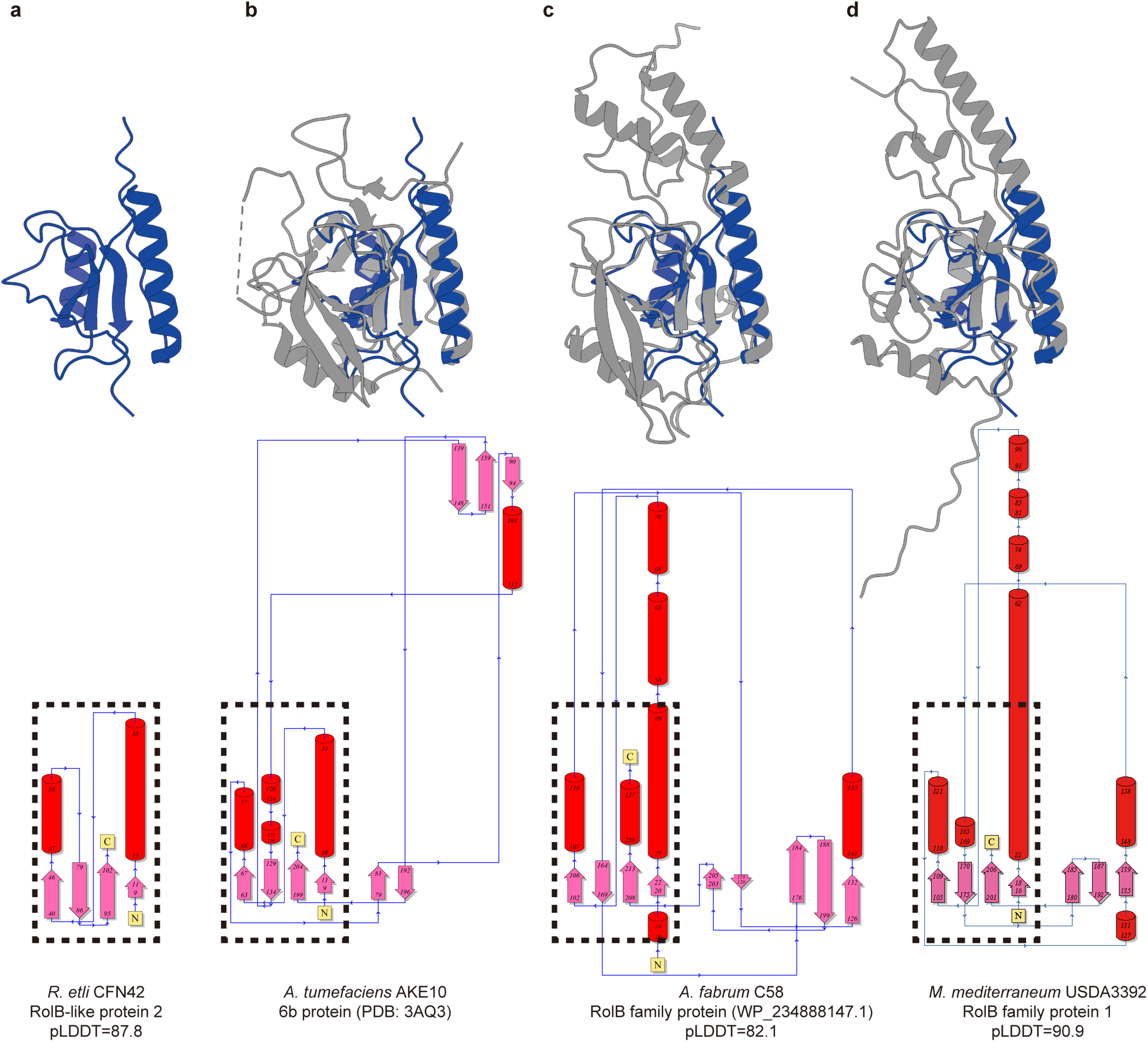
Structural comparison of RolB-like and RolB family proteins. **a**, Top, AlphaFold2-predicted structure of RolB-like protein 2 from *Rhizobium etli* CFN42, overlaid with the structures shown in **b**–**d**. Bottom, topology diagram of the same RolB-like protein; regions corresponding to structurally similar segments in **b**–**d** are indicated by dashed rectangles. **b**, Crystal structure of the *Agrobacterium* 6b protein (PDB 3AQ3). **c**, AlphaFold2-predicted structure of a RolB family protein (WP_234888147.1) from *Agrobacterium fabrum* C58. **d**, AlphaFold2-predicted structure of a RolB family protein encoded by a mini-T-DNA of *Mesorhizobium mediterraneum* USDA3392 analysed in this study.

**Extended Data Fig. 2.**
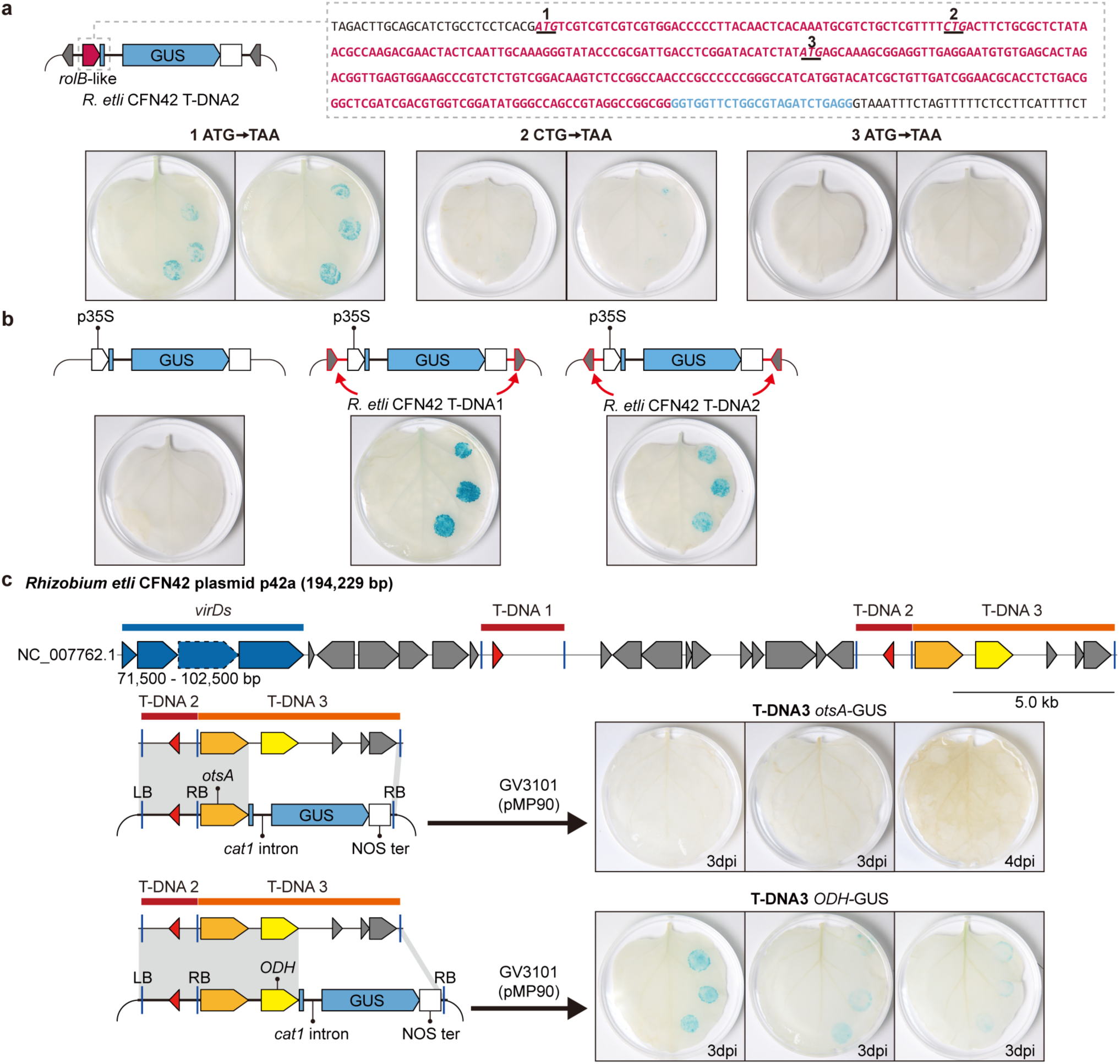
Additional analyses supporting Fig. 3. **a**, The translation start site of the *rolB*-like gene encoded by *Rhizobium etli* CFN42 T-DNA2 may not correspond to an ATG codon. To identify the likely start codon, candidate sites were individually mutated to TAA in the GUS-fusion construct and assessed by leaf infiltration. Candidate positions are labelled 1–3; position 2 is the most likely start site based on sequence homology (Fig. 3c), whereas positions 1 and 3 correspond to the nearest upstream and downstream ATG codons, respectively. **b**, To confirm that GUS activity reflects T-DNA delivery mediated by T-DNA border sequences, a binary vector carrying a p35S-GUS cassette but lacking T-DNA borders was tested as a negative control, alongside two otherwise identical constructs containing the T-DNA border sequences of *R. etli* CFN42 T-DNA1 or T-DNA2. **c**, The two cargo genes of *R. etli* CFN42 T-DNA3, which lies adjacent to T-DNA2, were also tested for delivery and expression by *Agrobacterium*-mediated leaf infiltration of *Nicotiana benthamiana*, as in Fig. 3c. Genetic organisation of the mini-T-DNAs is shown above; the left-hand schematic indicates the regions cloned into binary vectors, connected by shaded grey areas. *otsA*, trehalose-6-phosphate synthase gene; *ODH*, opine dehydrogenase family protein gene. dpi, days post-infiltration.

**Extended Data Fig. 3.**
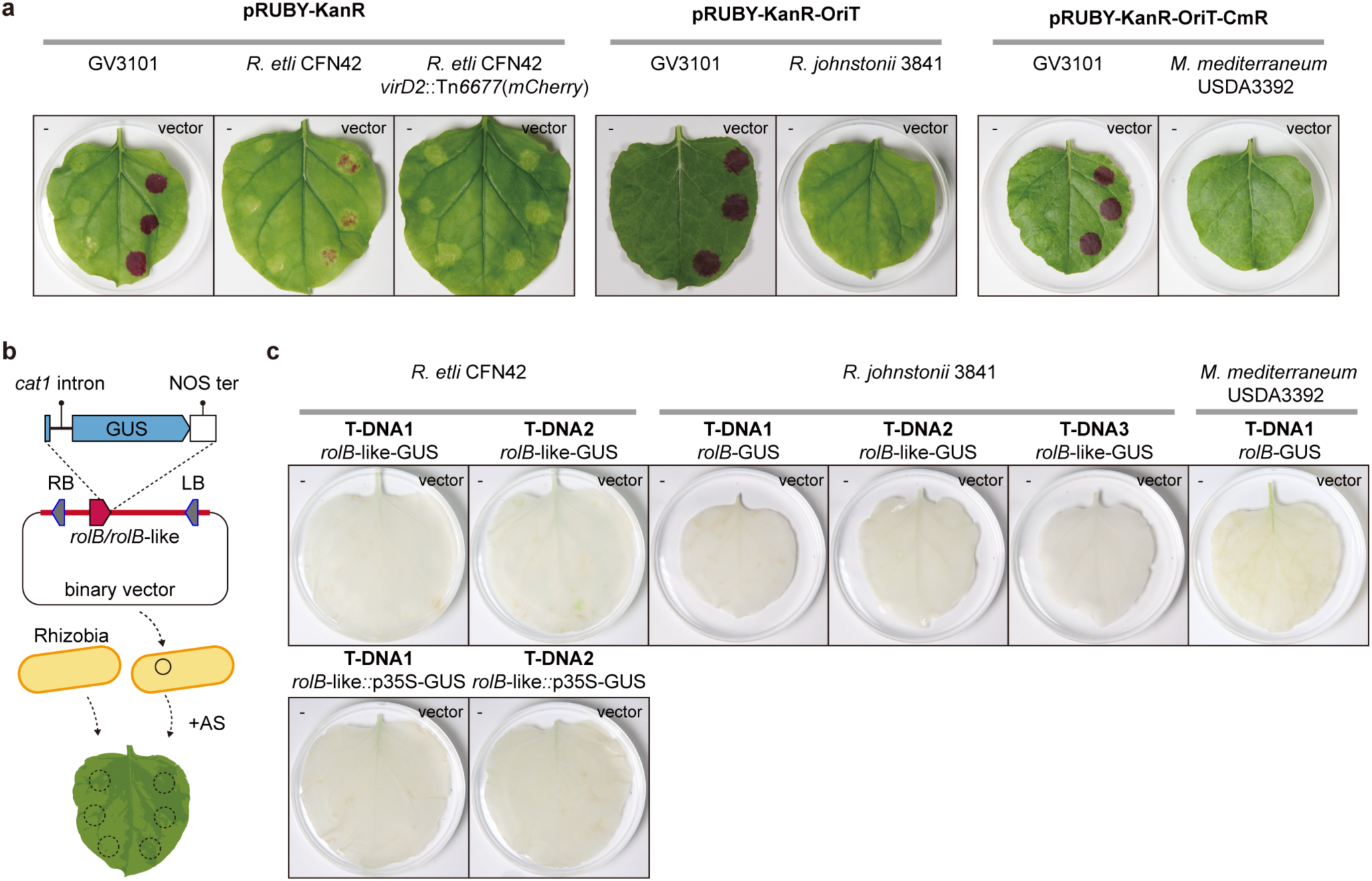
Testing the ability of the rhizobial strains to mediate T-DNA transfer into *Nicotiana benthamiana*. **a**, Three rhizobial strains carrying native T-DNAs were tested for their ability to transform *Nicotiana benthamiana* leaves. Constructs carry a *RUBY* reporter cassette, and modified variants tailored for each rhizobial strain were assayed, with *Agrobacterium fabrum* GV3101(pMP90) included as a positive control. Left, *Rhizobium etli* CFN42 and its *virD2*-disrupted mutant; middle, *Rhizobium johnstonii* 3841; right, *Mesorhizobium mediterraneum* USDA3392. **b,** Schematic illustration of the experimental setup. AS, acetosyringone; LB, T-DNA left border; RB, T-DNA right border. **c,** Assessment of the transferability of six T-DNAs and the expression of their *rolB/rolB*-like genes to *Nicotiana benthamiana* leaves by their native rhizobial hosts. T-DNAs of *Rhizobium etli* CFN42 are also tested with p35S in place of *rolB*-like genes.

**Extended Data Fig. 4.**
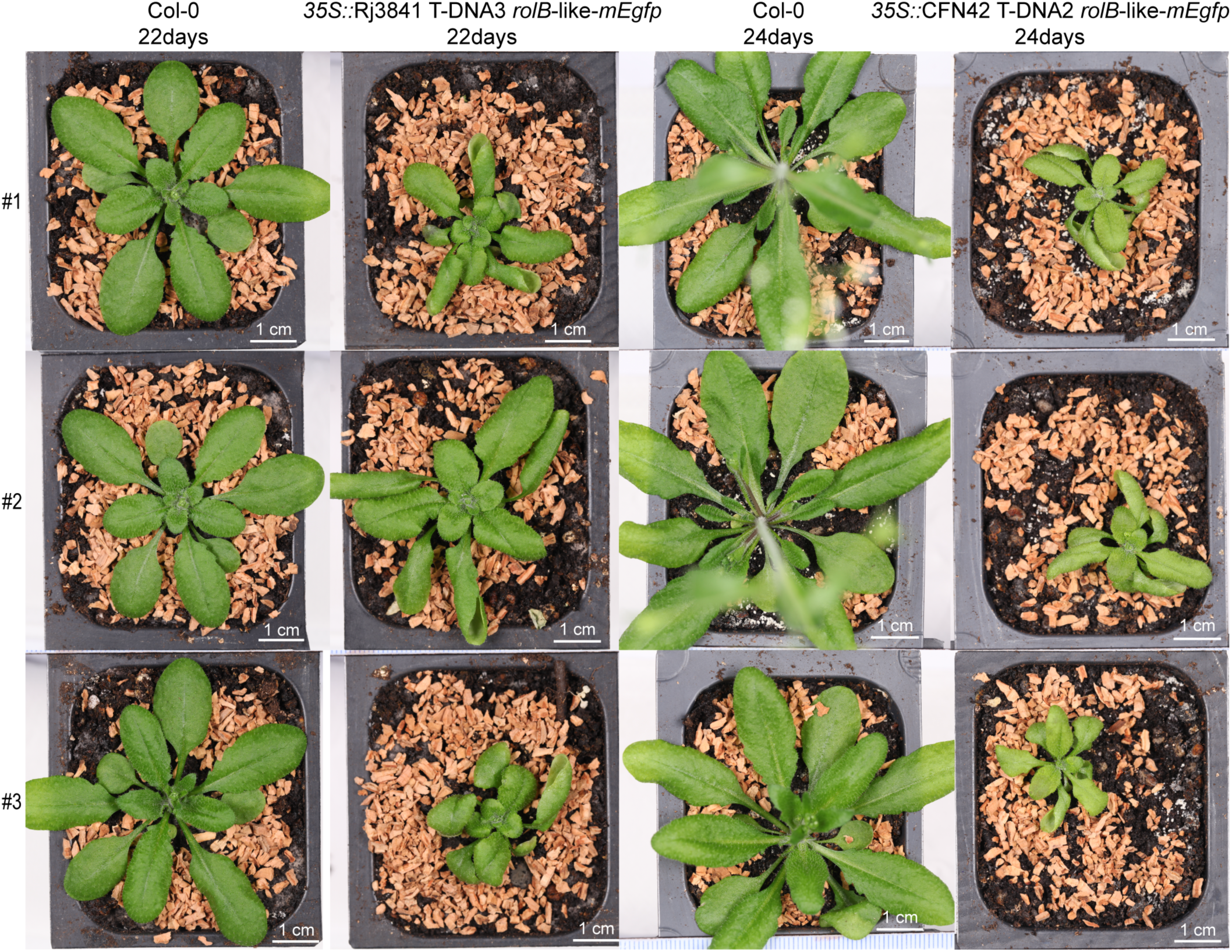
Transgenic *Arabidopsis thaliana* expressing rhizobial RolB-like-mEGFP fusions display leaf epinasty. Transgenic *A. thaliana* T3 lines expressing RolB-like-mEGFP from *Rhizobium johnstonii* 3841 (Rj3841 T-DNA3) or *Rhizobium etli* CFN42 (CFN42 T-DNA2) under the CaMV 35S promoter exhibited leaf epinasty and stunted growth. Plants were photographed 22-24 days after sowing, alongside wild-type *A. thaliana* Col-0 grown under the same conditions and at the same time. For each construct, one representative of three independently generated transgenic lines is shown; GFP fluorescence confirmed expression in leaves. The transgenic lines were not confirmed to be homozygous or to carry single-locus insertions.

**Extended Data Fig. 5.**
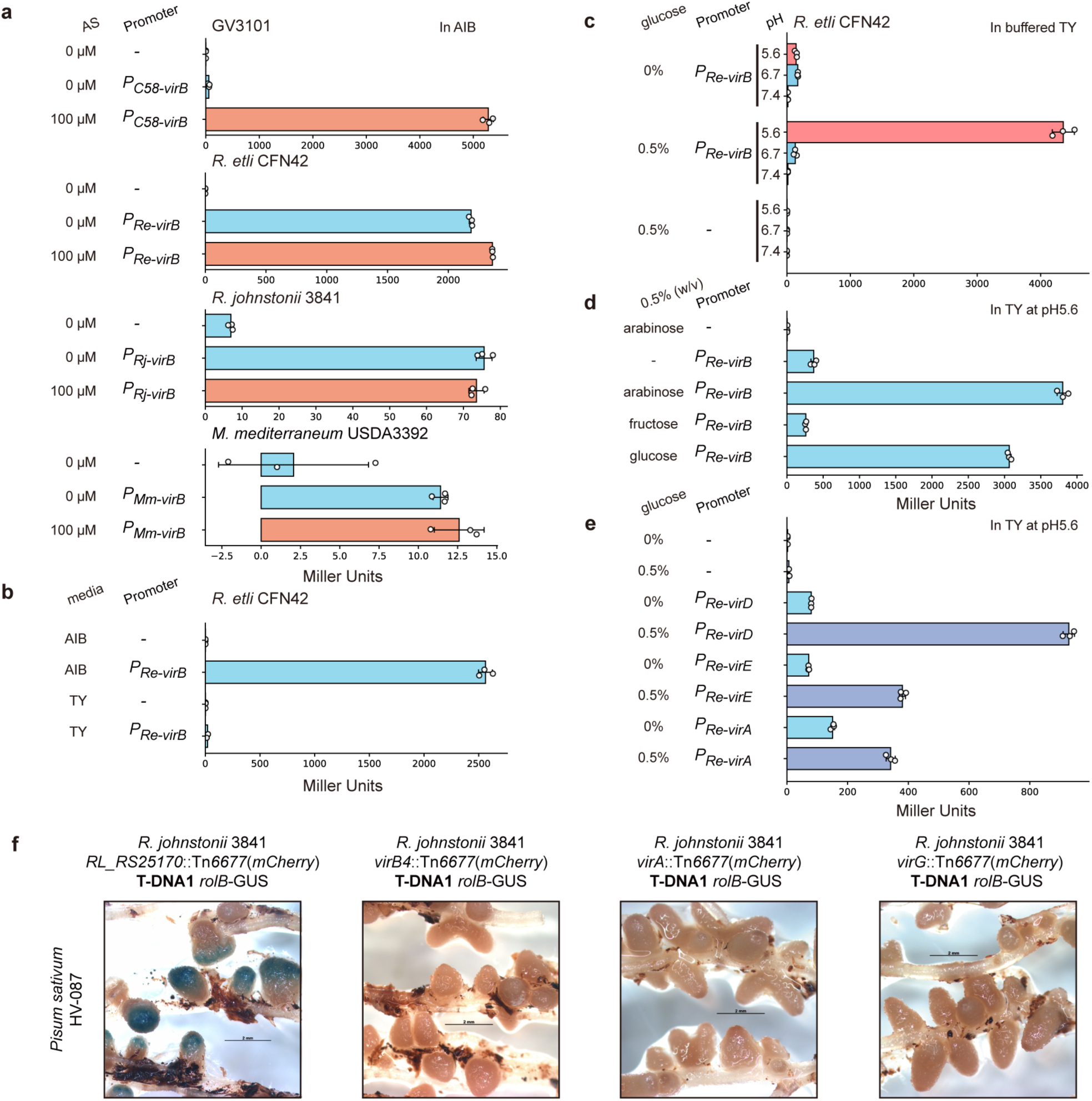
Responses of agrobacterial and rhizobial *vir* promoters to acetosyringone and culture conditions. **a**, Acetosyringone (AS) induction of *virB* promoters from *A. fabrum* GV3101(pMP90) and three rhizobial strains in *Agrobacterium*-inducing buffer (AIB). **b**, Activity of the *R. etli* CFN42 *virB* promoter in AIB versus TY medium. **c**, Effects of pH and glucose on *R. etli* CFN42 *virB* promoter activity in buffered TY medium. **d**, Under pH 5.6, arabinose and glucose, but not fructose, activate *R. etli* CFN42 *virB* promoter activity in TY medium. **e,** Effects of glucose on *R. etli* CFN42 *virD*, *virE*, and *virA* promoter activities in TY medium at pH 5.6. **f**, *R. johnstonii* 3841 carrying disruptions of *virB4*, *virA*, *virG* or an integrase gene (RL_RS25170 as a control), all located on pRL7, were tested for their ability to mediate T-DNA transfer into root nodules. Experimental conditions were as in Fig. 4. Tn*6677*(mCherry), a CRISPR-guided transposon carrying a constitutive mCherry reporter. Scale bars, 2 mm.

**Extended Data Fig. 6.**
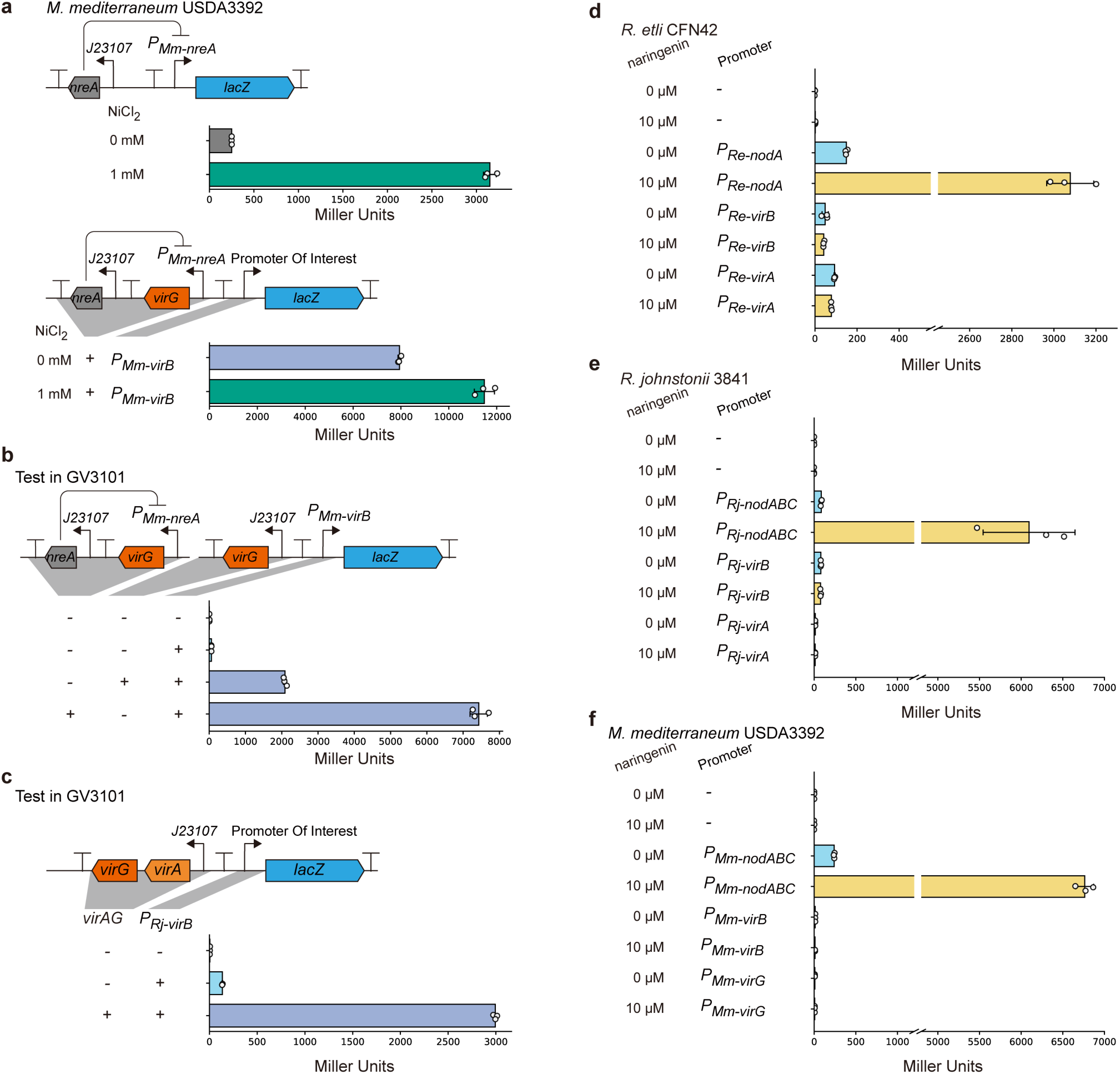
Effects of rhizobial VirG and VirAG expression and naringenin on *vir* promoter activity. **a**, To mitigate the growth defect of *M. mediterraneum* USDA3392 caused by VirG overexpression, a Ni²⁺-inducible expression system derived from *M. mediterraneum* USDA3392 was evaluated. This system relies on the repressor NreA, which dissociated from its cognate promoter (P*_Mm_*_-_*_nreA_*) upon binding Ni²⁺ ions. Placement of *M. mediterraneum* USDA3392 *virG* under NreA-mediated repression reduced the growth defect. Leaky VirG expression in the absence of Ni²⁺ was sufficient to induce the *virB* promoter strongly. **b,** Induction of the *virB* promoter by *M. mediterraneum* USDA3392 VirG is recapitulated in *A. fabrum* GV3101(pMP90). **c,** Induction of the *virB* promoter by *R. johnstonii* 3841 VirAG is likewise observed in *A. fabrum* GV3101(pMP90). **d**, Effect of 10 μM naringenin on the *R. etli* CFN42 *nodA*, *virB*, and *virA* promoters. **e**, Effect of 10 μM naringenin on the *R. johnstonii* 3841 *nodABC*, *virB*, and *virA* promoters. **f**, Effect of 10 μM naringenin on the *M. mediterraneum* USDA3392 *nodABC*, *virB*, and *virG* promoters.

**Extended Data Fig. 7.**
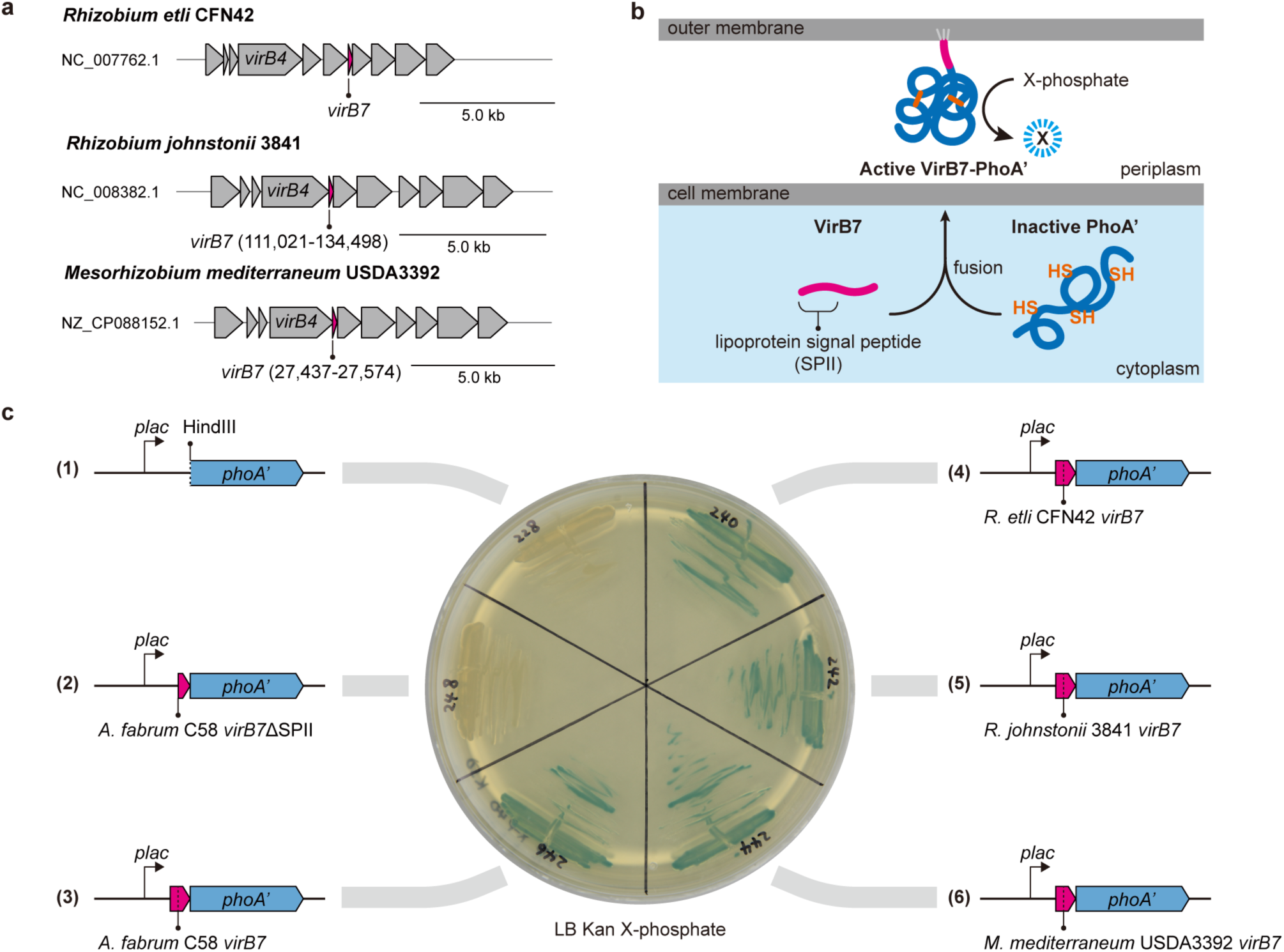
Putative VirB7 proteins from *Mesorhizobium mediterraneum* USDA3392 and *Rhizobium johnstonii* 3841 encode functional export signals. **a**, Schematic of the *virB* operons from the rhizobial strains used in this study. The *virB* operon of *Rhizobium etli* CFN42 resembles the canonical *Agrobacterium* organisation, whereas those of *M. mediterraneum* USDA3392 and *R. johnstonii* 3841 show low sequence identity to *Agrobacterium* and distinct architectures, in which the putative *virB7* gene lies between *virB4* and *virB5*. **b**, Schematic of the protein-export assay. *Escherichia coli* alkaline phosphatase (PhoA) requires an oxidative environment to form disulfide bonds and fold correctly, and is therefore active only when exported to the periplasm. PhoA lacking its N-terminal secretion signal (PhoA′) was fused to the putative VirB7 proteins, which are predicted to contain a lipoprotein signal peptide recognised by signal peptidase II (SPII); export of the fusion, and hence PhoA activity, reports whether these VirB7 homologues encode functional export signals. Active PhoA converts X-phosphate (5-bromo-4-chloro-3-indolyl phosphate) to a blue product. **c**, PhoA assay of the putative rhizobial VirB7 proteins (sections 4–6 of the plate). VirB7 from *Agrobacterium fabrum* C58 and its signal-peptide-deleted derivative served as positive and negative controls, respectively (sections 2 and 3), and PhoA′ alone as a further negative control (section 1).

**Extended Data Fig. 8.**
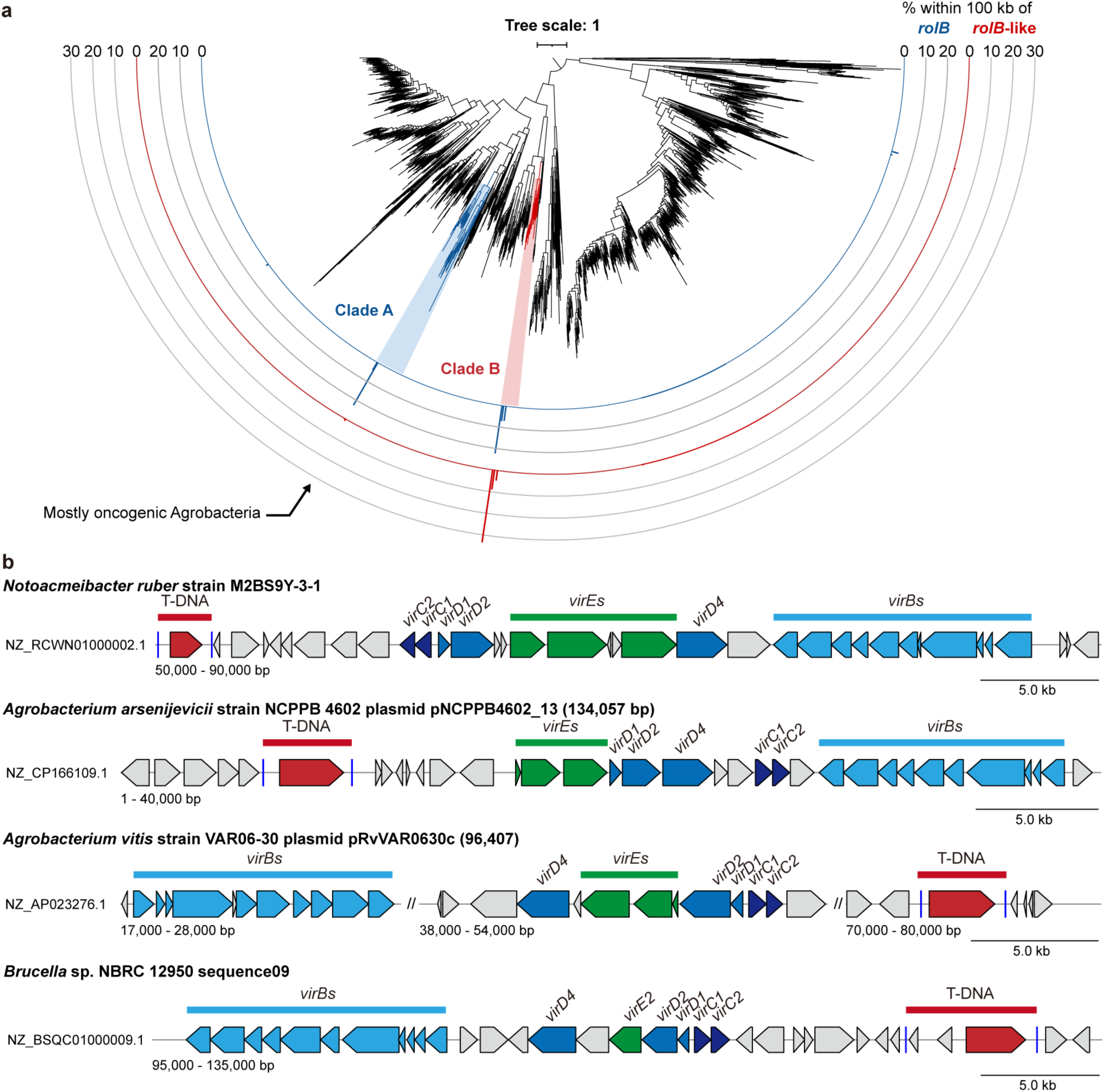
Phylogeny of alphaproteobacterial VirB4 and examples of putative T-DNA transfer systems in endophytes. **a**, Phylogenetic tree of alphaproteobacterial VirB4 proteins, constructed from representative sequences of clusters sharing >80% amino acid identity. For each cluster, the proportion of genomes with a *rolB* or *rolB*-like gene within 100 kb of *virB4* is shown as a bar chart. Two distantly related VirB4 clades co-localise with *rolB* or *rolB*-like genes: the clade containing most VirB4 proteins from oncogenic Ti and Ri plasmids (clade A) is indicated, and the other is clade B (as defined in the main text). Coloured clades are shown at higher resolution in Fig. 6. **b**, Genetic organisation of putative T-DNA transfer systems from representative endophytic and other non-oncogenic, non-rhizobial bacteria (*Notoacmeibacter ruber* M2BS9Y-3-1, *Allorhizobium vitis* VAR06-30, *Agrobacterium arsenijevicii* NCPPB 4602, and *Brucella* sp. NBRC 12950), corresponding to the clades highlighted in Fig. 6. Homologues of *Agrobacterium* T-DNA transfer genes and candidate T-DNA regions are indicated.

**Extended Data Fig. 9.**
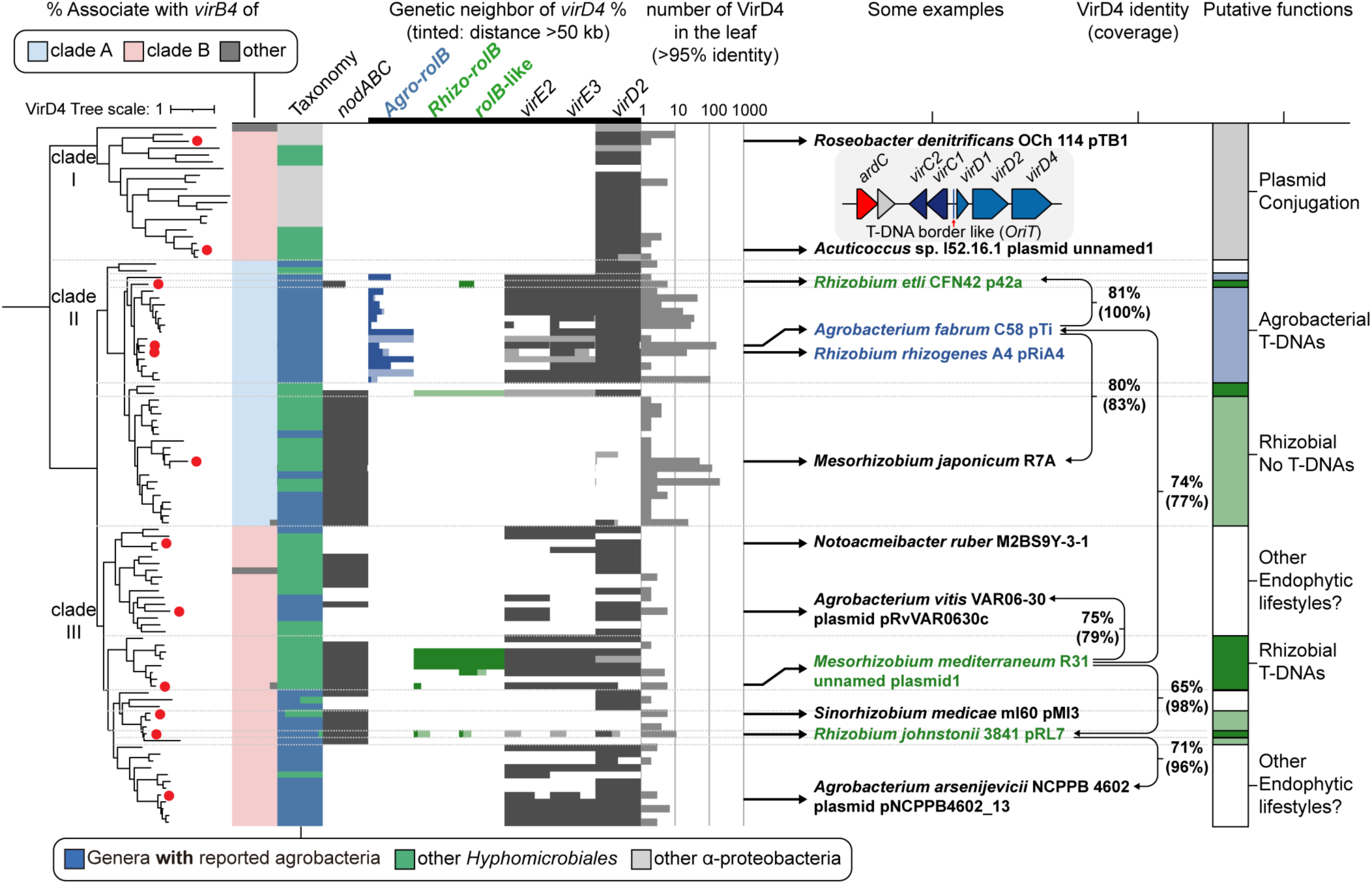
Phylogenetic analysis of alphaproteobacterial VirD4 proteins associated with agrobacterial T-DNA transfer systems. A phylogenetic tree of alphaproteobacterial VirD4 proteins related to agrobacterial VirD4, constructed from representative sequences of clusters sharing >95% amino acid identity. For each cluster, the proportion of *virD4* genes co-localising with clade A or clade B *virB4* loci, as defined in Extended Data Fig. 8a, is shown in two colours, and genus-level taxonomic composition is colour-coded according to NCBI taxonomy. The proportion of clusters whose genomes contain a *nodABC* gene cluster is shown in dark grey. The presence of selected genes on the same contig as *virD4* is indicated similarly, with light grey denoting genes located >50 kb from *virD4*. *rolB* genes are separated into two groups corresponding to the clades primarily associated with rhizobia or with agrobacteria, as defined in Fig. 1c. Cluster sizes are shown as bar charts on a logarithmic scale. VirD4 proteins are indicated for the rhizobial strains analysed in this study and for the well-characterized systems of *Agrobacterium fabrum* C58 (pTiC58), *Rhizobium rhizogenes* A4 (pRiA4), and *Mesorhizobium japonicum* R7A; selected VirD4 proteins from putative endophytic T-DNA transfer systems that are neither oncogenic nor rhizobial are also labelled. Pairwise identities between selected VirD4 proteins are shown as connecting lines, and putative functions of the T4SS subclades are indicated at right. The tree was rooted with a putative plasmid-conjugation VirD4 outgroup, chosen from a broader phylogeny of alphaproteobacterial VirD4 proteins built from representative sequences of clusters sharing >80% amino acid identity. *ardC*, anti-restriction protein ArdC, commonly associated with plasmid-conjugation genes.

## Supplementary information list

**Supplementary Table 1 | Bacterial strains and plants used in this study**

**Supplementary Table 2 | Plasmids used in this study**

